# The role of advective transport mechanism in continental-scale transport of fungal plant epidemics

**DOI:** 10.1101/2020.11.20.392035

**Authors:** Nolan Elauria, Taeah Truong, Kush Upadhyay, Oleg Kogan

## Abstract

In examining the continental-scale plant pathogen spread, we focus on the competition between the short-range stochastic hopping within the atmospheric boundary layer, and the laminar advection by the currents in the free troposphere. The latter is typically ignored, since it is assumed that the population of spores which have reached the troposphere is small, and the fraction of the remaining spores that survived the subsequent journey is negligible due to ultraviolet light and frigid temperatures. However, we claim that it is in fact a crucial mechanism for continental-scale spread. We argue that free tropospheric currents can not be ignored, even as the probability for spores to reach them and to survive within them approaches zero. In other words, models that neglect tropospheric advection are fragile – their predictions change qualitatively if this alternative transport channel becomes accessible – even when the rate at which spores actually make use of this transport channel approaches zero.

## 1 Motivation

The year 1923 was particularly bad for wheat growers in the United States. Farmers from Texas to Minnesota experienced a wind-driven epidemic of a fungal pathogen known as Wheat Rust, advancing up to 54 km/day [1]. A global pandemic remains a possibility in our times. Recently, a hypervirulent race Ug99 of the wheat stem rust fungus *(Puccinia graminis* f. sp. *tritici*) has generated considerable attention [2, 3, 4]. Although human activity, such as transport of spores on clothing or vehicles, are undeniably important transport mechanisms in the modern world, previous studies have also documented or supported atmospheric long distance transport of plant pathogens [2], [5]-[11], [12].

Atmospheric transport of passive scalar – fungal spores, aerosols, and dust – involves multiple transport mechanisms [14], [15]. The lowest layer of the atmosphere, known as the atmospheric boundary layer (ABL), is where most of the atmospheric turbulence is localized [16, 17]. The thickness of the ABL varies in time and space, but a typical value in wheat-growing mid-latitudes is 1-3 km during daytime [16]. Therefore, transport of spores over scales below several kilometers would be dominated by the aerodynamics of the ABL, and driven by short, random, turbulence-driven hops.

Because spore transport at scales below several kilometers is predominantly stochastic, the predictions of spread over these distances has been based primarily on models for dynamics of probabilities or densities [18]. In particular, kernel-based models received intense attention in recent years [20, 21, 22, 23] and [24] (the latter source concerns seed dispersal). In their essence, kernel-based models represent random walk on multiple scales; they could also be rightfully called “reaction-dispersal” models. The popularity of kernel-based models is probably due to the fact that detailed data on dispersal is only available on a kilometer scale (“plot scale”) or below [18, 20, 25, 26], and typically comes in the form of dispersal kernels measured from release and collection experiments.

Although ABL remains a possible mechanism of transport over distances much longer than plot scale – for instance, over entire continents – it is inefficient at such scales, because turbulent vortices of the ABL tend to return the passive scalar to the ground over distances comparable to its own thickness. This inefficiency of the ABL for transporting particles over continental-scale distances is well known. For example – the flows in the Saharan Air Layer, which is located above the marine layer, are the primary mechanisms of dust transport between Africa and Europe or even North America [27]-[28]. Fungi and spores are also transported by this mechanism [29]. Pollutants from Asia are also reported to reach North America by means of long-range winds above the ABL [30].

Transport facilitated by the free atmosphere (FA) that lies above the ABL is less random than the boundary layer driven transport. Thus, models based on dispersive kernels, which inherently describe a random method of transport, become inapplicable. At the same time, it is known that spores of certain pathogens are resistant to ultraviolet radiation and are able to withstand many hours in the air under the cloudy conditions [1], [54], [55], [?] (and references within – see Chapter 6). This suggests to us that FA could emerge as a relevant transport mechanism over continental scales.

The analysis of the role of advective method of transport – in contrast to a random walk method described by kernel-based models – has been proposed before. Examples of such advection-based analyses include Gaussian puff model [1], [11], [12], [31], and Lagrangian stochastic simulations [26], [32]. The former describes the dynamics of a plume of spores – including the effect of ultraviolet radiation on survivability of propagules and dilution due to the stretching and spreading out of the plume. It makes a prediction about the rate of deposition from such a plume unto the ground.

One question that has not been examined, with the exception of our previous work [33], is the competition between the two mechanisms of transport – the random shortrange hopping transport mechanism, versus the non-random advective transport mechanism. To summarize, the first mechanism bears the advantages of being efficient for short-range transport and not requiring spores to spend long times exposed to UV radiation, but suffers from the disadvantage of being inefficient over distances that are much greater than several kilometers. The advantages of the second mechanism are its speed and efficiency in long range, but the drawbacks are: (i) inaccessibility – probability of spores to reach troposphere is small and (ii) UV suppression – spores that do reach it end up being exposed to UV radiation for significant periods of time because of a longer residence time there, as we will discuss in the main text. This naturally brings up the following question – when both methods of transport are available in a model, which one will be dominant for continental-scale spread? In other words, how does the availability of the advective transport channel modify the spatiotemporal dynamics of the spore density and disease density during the spread? Would opening of this second, non-random transport channel change the invasion dynamics, or would the availability of the second transport channel – as efficient as it is from mechanical perspective – not significantly affect the spatiotemporal dynamics of invasion fronts because of inaccessibility and UV suppression? This is the question that we address in this paper.

The first step in this investigation, focusing specifically on these two competing effects, was taken by one of us recently [33]. The random mechanism of transport was modeled as diffusion, which competed with the non-random advective mechanism. The random mechanism acts on spores close to the ground and within the ABL, while the non-random one acts in the higher atmosphere. Because each mechanism operates at a different height, advection could not be removed by a simple change of a reference frame. This stands in contrast to a simple Fisher-Kolmogorov model with an additive advection term – here transforming into a frame that is co-moving with the advective flow would get rid of the the advective term. The key finding of this paper was the following: opening up of the second, non-random advective transport qualitatively changes the spatiotemporal dynamics of invasion fronts. What is especially remarkable, is that this qualitative change to the dynamics takes place even when the probability of reaching the advective layer goes to zero (but is not equal to zero). The term “qualitative change” means the following – a model that does not include the advective mechanism will predict a different dependence of the front properties upon parameters, such as diffusion coefficient, than a model that does include the alternative advective transport channel. Front properties refer to speed and shape. In mathematical literature, a qualitative change in the nature of the solution upon an infinitesimal change of a parameter is known as a singular perturbation [34].

Our result directly disproves the assumption that free troposphere will not matter due to inaccessibility. Inclusion of a separate advective channel of transport results in a qualitative and significant change in spatiotemporal dynamics – even if the fraction of spores that take advantage of this channel is infinitesimal. We will call this “fragility effect” – because it indicates that a diffusive model is fragile with respect to opening up of an alternative competing transport channel.

Fragility was proven and explored at great depth in [33] within a context of a simplified model, where the spore death from UV exposure and latency for producing new spores was not included. Thus, while it addressed the issue of inaccessibility, it did not tackle UV suppression. But most importantly, this basic model over-predicted the invasion speed. The present work addresses these issues by including the missing biological processes in the model. The resulting extended model depends on the following additional parameters: the rate at which spores infect host plants, the rate at which airborne spores die due to UV, and the latent time required for new fungi to start producing new uredospores. The resulting extended model is capable of making more realistic predictions of the front speed.

An important question is whether the fragility persists under the said extension of the basic model, where the missing processes are included. Remarkably, we find that fragility effect does not go away. It remains true that an arbitrarily small, but finite coupling to the advective transport by the FA qualitatively changes the spatiotemporal dynamics of invasion waves. This is our main result of the paper – fragility effect is robust with respect to considered extensions of the model.

In summary, opening up of the second, non-random advective transport channel qualitatively changes the spatiotemporal dynamics of invasion waves, even as the probability of reaching the advective layer goes to zero. The effect is found to be robust with respect to variations of the model, where increasingly more complex elements of biological realism are taken into account. It is important to know that we are not attempting to be meteorologically rigorous. Our study was designed specifically to address the question of the competition between a random and a non-random mechanism of transport. Therefore, it was not necessary to stick to meteorological rigor. We merely needed a simple model which accommodates the competition between random and non-random transport mechanisms.

We hope that our finding of robustness of fragility will lead to re-examining of the role of free troposphere in understanding of continental-scale spread of fungal crop epidemics, and of control strategies. We also hope that it will inspire new simulations of continental-scale transport, and potentially, long-range observations involving the use of genetic markers.

The paper is organized as follows. In Section 2 we outline the findings from our previous work, and explains fragility. Section 3 introduces new biological realism and demonstrates robustness of fragility with respect to these changes. The key result is presented in Fig. 9. We reflect on the new findings in Section 4.

## 2 Summary of previous findings

### 2.1 Model philosophy

Dispersal within ABL is a result of two transport processes working together: vertical diffusion due to turbulent eddies, and horizontal advection by mean winds. As a particle performs random walk in the vertical direction, it is at the same time advected downstream. The net effect is a heavy-tailed distribution of landing locations – also known as contact distribution in the literature, or simply dispersal kernel. Theoretical predictions of dispersal kernels based on this picture have been developed in the past [35] (and references therein), [36].

In the absence of a horizontal component of random walk, the dispersal kernel would be zero in the upwind direction of mean winds would be strictly zero. Thus, it would be skewed strictly in the downwind direction of mean winds in the ABL. However, the random walk has a horizontal component as well, which gives this kernel a tail in the upwind direction, but its peak would be shifted in the downwind direction. Therefore, the effect of such dispersal can be represented by a combination of a kernel with a peak at zero, plus an advective velocity.

It is also known that eddy diffusivity tapers off to zero at the top of the ABL [37]-[39]. In the free atmosphere that lies above the inversion layer, eddy diffusivity can be neglected, although there are notable exceptions, such as clouds, thunderstorms, etc.

The dynamics of penetration through inversion layer appears to be a topic of open debate in the atmospheric science community. Penetrating the inversion layer is not easy, and most of the material that crosses it does so in the morning and at night, when the thickness of the ABL changes rapidly. In the morning hours, as the ABL grows, droplet-like structures of the inversion layer are left behind, providing a potential mechanism for particles to cross from the FA into ABL [40]. In the evening hours, when the ABL shrinks, some material may be left behind in the residual layer, and eventually end up in the FA **cite**. This is a potential mechanism for particles to cross from ABL to FA. During the day, however, and in the presence of a developed inversion layer, the probability for crossing is small in the sense that the time scale to cross the inversion is much greater than the time scale of the largest ABL eddy. Therefore, if particles end up in the FA, they are likely to stay there for a time that is much longer than the timescale of the largest eddy in the ABL below – there is a natural timescale separation.

### 2.2 Constructing the model

The discussion above leads to the following picture. The ABL transports spores by a combination of advection by mean winds and horizontal dispersion. This random horizontal dispersion results primarily from a combination of vertical turbulent dispersion and horizontal mean winds in the ABL [35] or [36], although there is also some effect due to horizontal turbulent dispersion as well. ABL also gives rise to the vertical random motion, which moves particles to and from the boundary with the free atmosphere. The turnover time is known to be on the order of 1/2 hour. The free atmosphere is, to an approximation, purely advective. The residence time of particles in the FA is much longer than the 1/2 hour vertical turnover time.

Because of the differences in the residence time, we propose a two-layer model, where the separation between the layers is made by the criterion of residence time. The ground layer (GL) represents the ground and the ABL. The residence time of particles in this layer is on the order of a 1/2 hour timescale of the largest eddies; particles typically return to the ground over this time scale. The mean wind velocity in this layer is *v_GL_*. The second layer represents the free atmosphere, where the residence time of particles is expected to be significantly longer due to the difficulty of penetrating through the inversion layer that separates it from the ABL. We will call it advective layer (AL), because advection is the only mechanism of transport there. The advective velocity in this layer is *v_AL_*, typically larger than *v_GL_* **cite**.

Because our main objective here is to study the competition between a random horizontal dispersive mechanism vs. the advective mechanism, for the purpose of this paper we will work with diffusion, rather than dispersion. The coefficient of turbulent eddy diffusion is *K* (the notation comes from what’s called “K-theory” in the field [26]).

Choosing to model the random walk by diffusion, rather than a heavy-tailed dispersion is not just a matter of model simplicity. It is well known that exponentially-decaying dispersal kernels give rise to an effective diffusion [41] on a length scale much greater than the scale of the decay of this exponential kernel. On the other hand, when dispersal kernels have powerlaw tails – which is the case with turbulence – one can not represent it by a diffusive term on any length scale. However, it is critical to keep in mind that diffusion is localized to the ABL. Thus, the largest size of turbulent eddies is comparable to the thickness of the ABL, which is at most several kilometers. Although it is possible for a particle to experience a horizontal transport by means of turbulent eddies over a much longer distance, this would constitute a rare event; a typical transported particle will return back to the ground over a length scale of several kilometers. Therefore, power law kernels have an exponential cutoff; power law dispersal kernels for turbulence-driven horizontal transport cross over to exponentially decaying ones over the length scale of several kilometers. This paper is concerned with horizontal transport over hundreds or even thousands of kilometers. Therefore, replacing dispersion by diffusion is completely justified.

The number density of spores in the advective and ground layers is *ρ*(**x**,*t*) and *σ*(**x**,*t*) respectively. The advective layer has an imposed velocity field **v**(**x**). There is a rate *α* > 0 at which spores switch from GL to AL (desorption), and a rate *β* > 0 at which spores switch from AL to GL (adsorption). The transport of particles from GL to AL is accommodated by vertical diffusivity, especially by the largest eddies, which bring the particles to the inversion layer. We are not deriving parameters *K, α*, and *β* from the details of the ABL dynamics; these are simply phenomenological parameters in our theory. However, we estimate these in Appendix A.

It was stated above that the dominant mechanisms of penetration through the inversion layer are active primarily in the morning and evening hours, so it might seem that *α* and *β* must be periodic, reflecting these diurnal cycles. In fact, we studied on the role of the temporal variation of these parameters, and found that the amplitude of the resulting variation of the front speed is actually minor for realistic values of *α* and *β*. This is discussed in Appendix D.

Finally, there’s the spore production. Spores produce other spores on the GL, and the growth rate is density dependent, given by some function *δf* (*σ*), where the characteristic growth rate *δ* is explicitly factored out of *f*. For example, for a logistic growth rate, 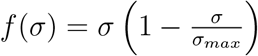, with *σ_max_* representing the carrying capacity. Thus, the basic model does not differentiate between spores (propagules) and fungus (disease), and effectively treats spores as a “self-reproducing dust”. The new extension of the model considered in Sections 3 distinguishes spores density from disease density.

We will work in one dimension. All together, the densities *ρ*(**x**,*t*) and *σ*(**x**,*t*) will evolve according to

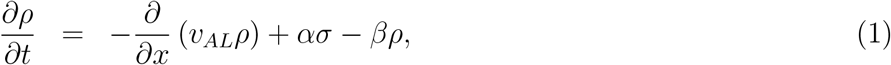

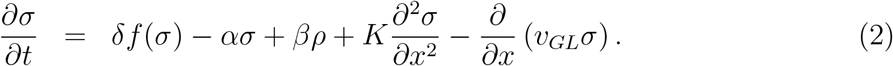

We will work with constant *v_GL_* and *v_AL_*. The model can be simplified by switching to a reference frame co-moving with velocity *v_GL_*. Then only the AL will contain the advective term

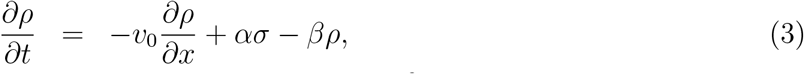

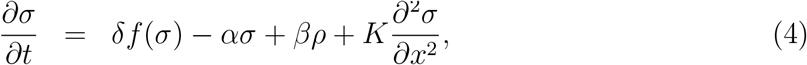

where *v*_0_ = *v_AL_* — *v_GL_*. The model can be summarized by the following schematic diagram.

There are five parameters in the model: *v*_0_, *α, β, δ, K*, which even at this simple level may seem unwieldy. However, they are not all independent. To see this, note that the model has a built-in natural unit of time, 1/*δ*, and a natural unit of velocity, *v*_0_. So, a natural unit of distance is *v*_0_/*δ*. By working with these units, rather than meters and seconds, we will be able to remove some parameters. Rescaling *x* by *v*_0_/*δ, t* by 1/*δ*, as well as *ρ* and *σ* by the carrying capacity *σ_max_*, we are left with only three parameters: 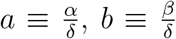 and 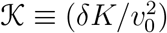. These units are natural for this problem, and in terms of them, the equations read

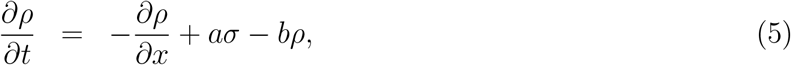

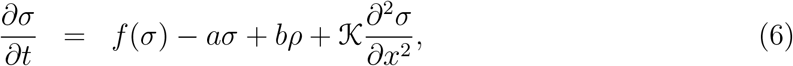

where we are still using letters *x, t, σ* and *ρ* to avoid cluttering the notation. If *f* (*σ*) is logistic growth function, it would read *σ*(1 – *σ*) in these rescaled units.

The model given by Eqs. (5)–(6) was thoroughly studied in [33]. It describe the average dynamics – they do not capture demographic stochasticity that results from finite-number fluctuations. We now summarize the key results.

### 2.3 Predictions the basic model

We now summarize the key results of this basic model; details can be found in [33]. It will prove to be very instructive to first describe the behavior in the regime when diffusion is set to zero. Thus, the diffusive channel of transport is turned off, and spores can move only by entering and exiting the advective layer. The model leads to one or two traveling density fronts, depending on the following criterion. If *aσ* < *f* (*σ*) for some 0 < *σ* < 1, there is a single front. On the other hand, if *aσ* > *f* (*σ*) for all 0 < *σ* < 1, there are two traveling fronts – the leading “upwind” front, and the trailing “downwind” front. For logistic growth, a single front happens for *a* < 1, i.e. desorption is slower than production of new spores, and two fronts happen for *a* > 1, i.e. desorption is faster than production of new spores. Both situations are demonstrated in Fig. 2.

**Figure 1:**
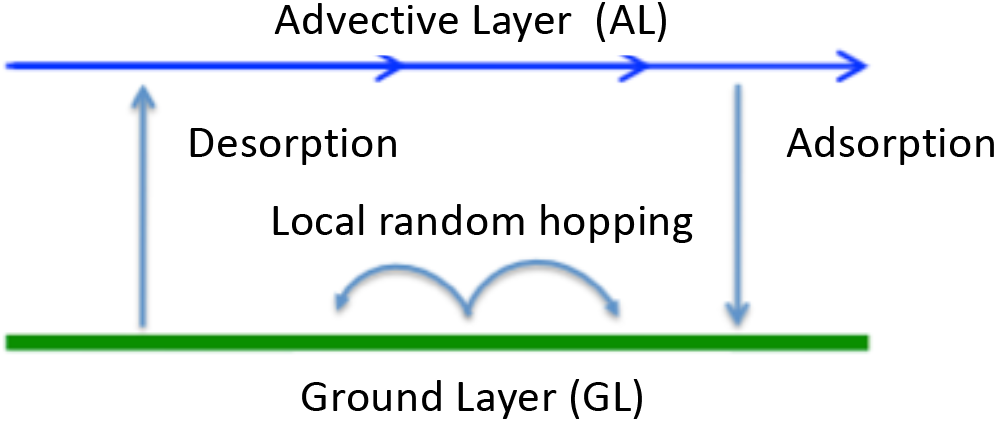
(Color online) Schematic of the basic model of [33].

**Figure 2:**
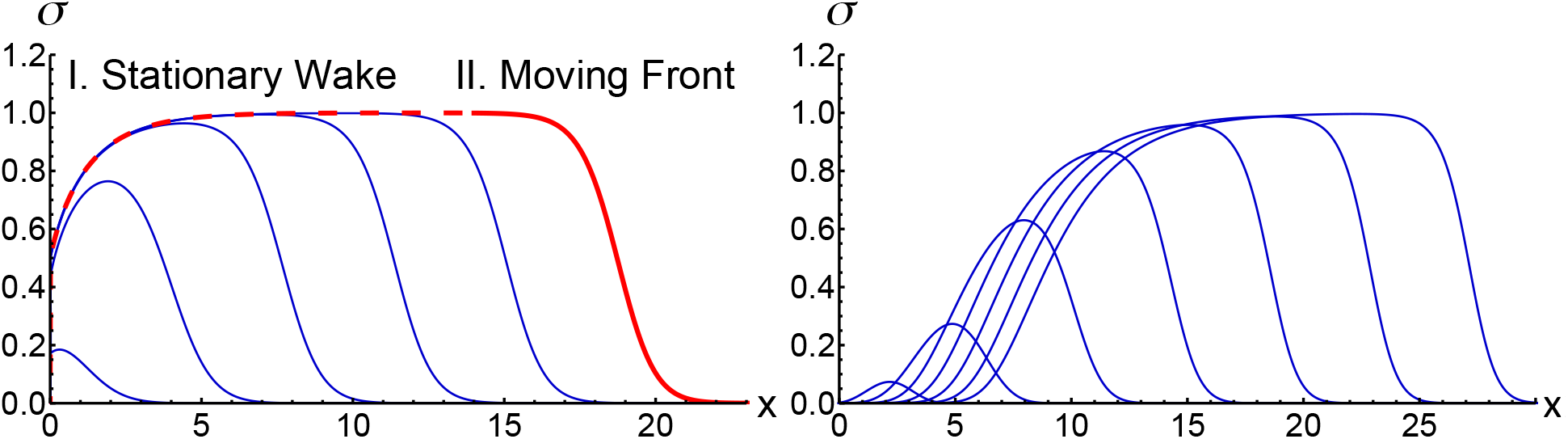
(Color online) Evolution of the GL profile from a point initial condition with a logistic growth model and no diffusion. Left: *a* = 0.5, *b* = 1 at *t* = 5, 10, 15, 20, 25 and 30. Right, *a* = 2, *b* =1, same ts. In both cases, the IC launches uniformly translating fronts. Early transients are not shown.

Such traveling density fronts are characterized by their width and speed. Here we focus on the speed, and study it as a function of parameters.

Fig. 3 summarizes results for a point initial condition in the GL. Continuous curves are analytical solutions, given by

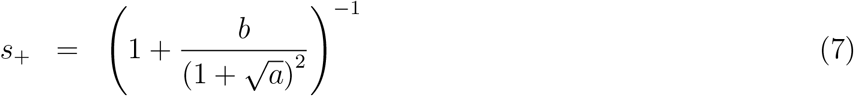

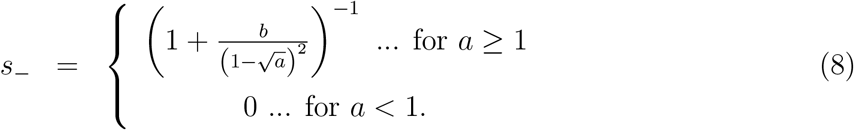

**Figure 3:**
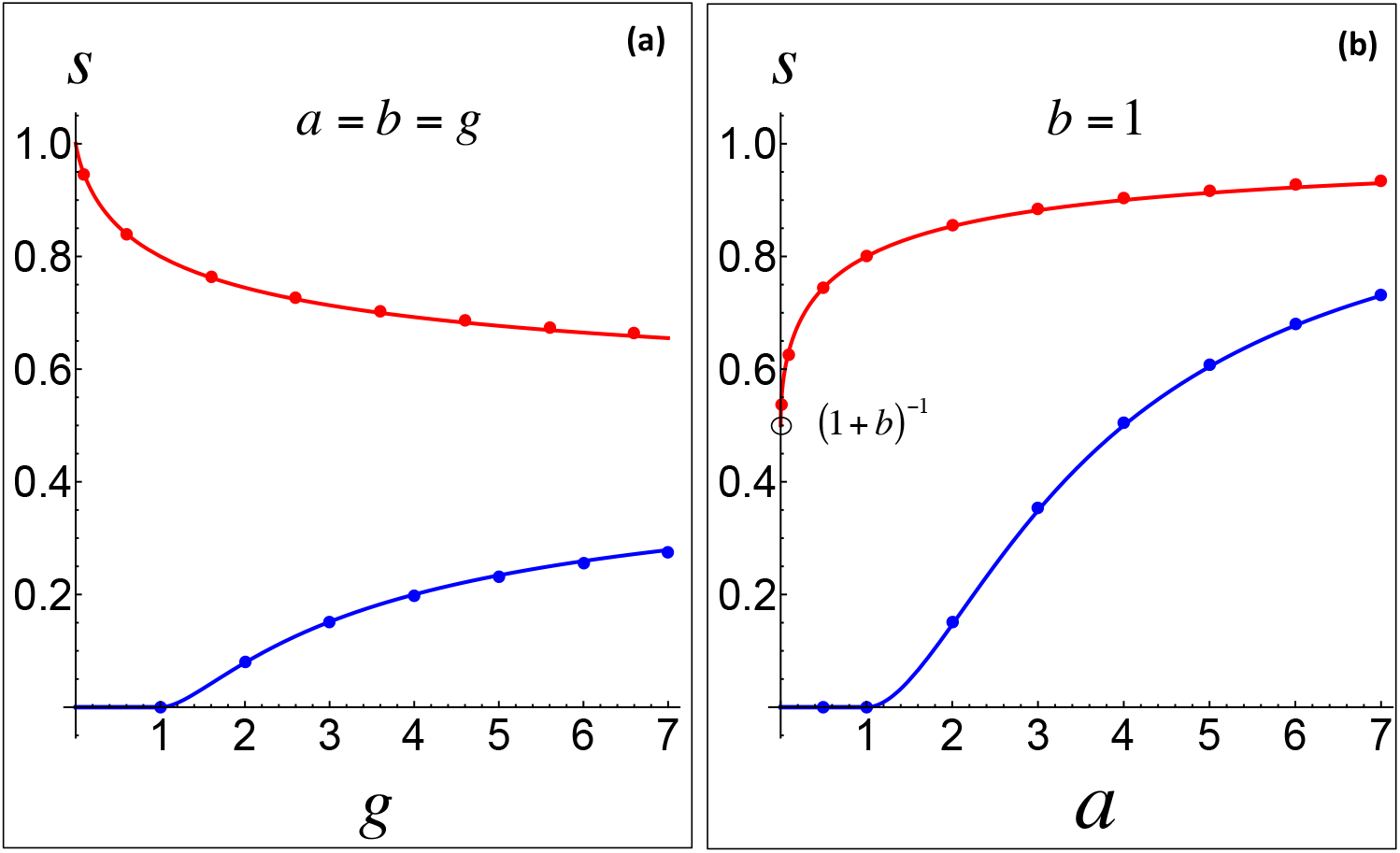
(Color online) (a): front speeds vs. the interlayer coupling *a* = *b* = *g* for a point initial condition in the GL. (b): front speeds vs. *a* at *b* =1. Here *s*_+_ → (1 + *b*)^-1^ as *a* → 0. The first two dots are at *a* = 0.01 and *a* = 0.1. Upper curve (red): downwind front speed *s*_+_, lower curve (blue): upwind front speed *s*_. Solid dots are from the numerical solution of Eqs. (5)–(6). Continuous curves are analytical solutions, given by Eqs. (7)–(8).

It demonstrates the key feature of the theory in the absence of diffusion: while at zero a, the front speeds will certainly be zero, the downwind front speed *s*_+_ remains finite as *a* approaches zero. These results apply for any growth function for which fronts are “pulled” [33], i.e. growth function for which the properties of the front (speed and decay rate) are governed by the leading edge, where the nonlinearities are not important [42]. Logistic growth function is of that type, because a theory based on the pulled front assumption match with numerical results of the model with a non-linear term (growth limiting term), see Fig. 3 and [33].

The prediction that a front speed goes to a non-zero value as the rate of entering the transport layer goes to zero is surprising. A parcel of mass that enters the AL – for however brief a period of time – will travel with the advective wind speed downstream, and because this is a continuum theory, there will always be mass present in the AL. So, the seeding process advances with the speed of the wind. The speed of the front is defined at a constant density contour, so in general it is less than the wind speed.

Consider, for example, the case when *b* is held fixed. As *a* → 0, the AL *density ρ* does go to zero, but the *speed* remains finite. At sufficiently small *a*, the particle density in the AL becomes so small that the continuum theory will break down. In [33], we provide a criterion for this to happen in, and conclude that for realistic atmospheric mixing conditions, this will never be a concern; the number of spores produced even on a single farm plot is so large, that a continuum approximation will always hold.

We can also understand this phenomenon from a different perspective. Consider ignoring the effect of desorption from the GL into the AL everywhere, except at the location of the source, i.e. ignore the ασ term in Eqs. (5)–(6). There is growth at the source. A fraction a of the amount produced there in time *dt* is transferred to the AL and carried downstream, while shedding mass to the GL. The evolution of *ρ* downstream from the source will echo the evolution at the source. For linearized equations, *ρ* is then given by a traveling exponential profile with a prefactor *α a*. Thus, the AL density itself goes to 0 as *a* goes to 0. The speed of this traveling exponential (measured at a constant contour, as usual) will be set by the speed of the advective layer. However, this speed will be less than 1 due to shedding – except when *b* also goes to zero, in which case the speed goes to 1. We can see this in the left panel of Fig. 3. The GL density *σ* will also be a traveling exponential, and will move with the same speed. The key message here is that even in the extreme case when desorption is completely turned off everywhere but at the source, the speed of the front will remain finite. The actual, full solution of *ρ* and *σ*, which includes both adsorption and desorption, is given in [33].

The phenomenon of the finite front speed in the limit of an infinitesimal exchange rate has been reported in other models of invasion dynamics. Lewis and Schmitz [43], for instance, considered a model of a population of individuals with two states – a diffusive state without reproduction, and an immobile state during which organisms reproduce. They also found that the invasion front speed approaches a finite value as the switching rate between the two states goes to zero. Another model was considered by Cook (see description in Ref. [44], Ch. 13) describing individuals who are always either stationary or diffusing, with each subpopulation reproducing according to a rate that depends on the total population. This model also exhibits a similar effect.

The finiteness of the leading front speed as the probability of reaching the AL goes to zero is at the very heart of everything else that follows. This result by itself is already suggestive that AL is crucial – that AL can not be ignored. However, this statement is only meaningful when this advective transport mechanism is compared to a competing transport mechanism. We will do that next, by adding diffusion to the GL, and demonstrating that AL remains crucial. That is – in the absence of AL, front speed has a different dependence on parameters than in its presence. Ignoring AL will give rise to qualitatively different predictions.

The situation that involves both transport mechanisms – advective and diffusive is summarized in Fig. 4 for downwind fronts. In the absence of advective transport channel (or mechanism), the model reduces to the Fisher-Kolmogorov-Petrovskiy-Piskunov (FKPP) model, with a well-known square root behavior of speed versus diffusion coefficient. The speed versus 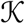 follows the “diffusion-only branch”. Opening up a competing diffusive trans-port channel causes a finite change in results – rather than a square root function, it is now an entirely different function. The speed vs. 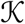 now follows a “coupled branch” consisting of solutions at non-zero *a* and *b*. For example, if *g* = 0.01, the new function is a constant up until 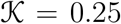, after which it follows the FKPP trend. Note that this drastic jump is actually largest as the coupling *g* goes to zero. This was already seen in the *a = b* panel of Fig. 3. As the coupling goes to infinity (pink curve), the jump is also finite, although not as large. This drastic change in results due to infinitesimal coupling to an advective transport channel is an example of fragility.

**Figure 4:**
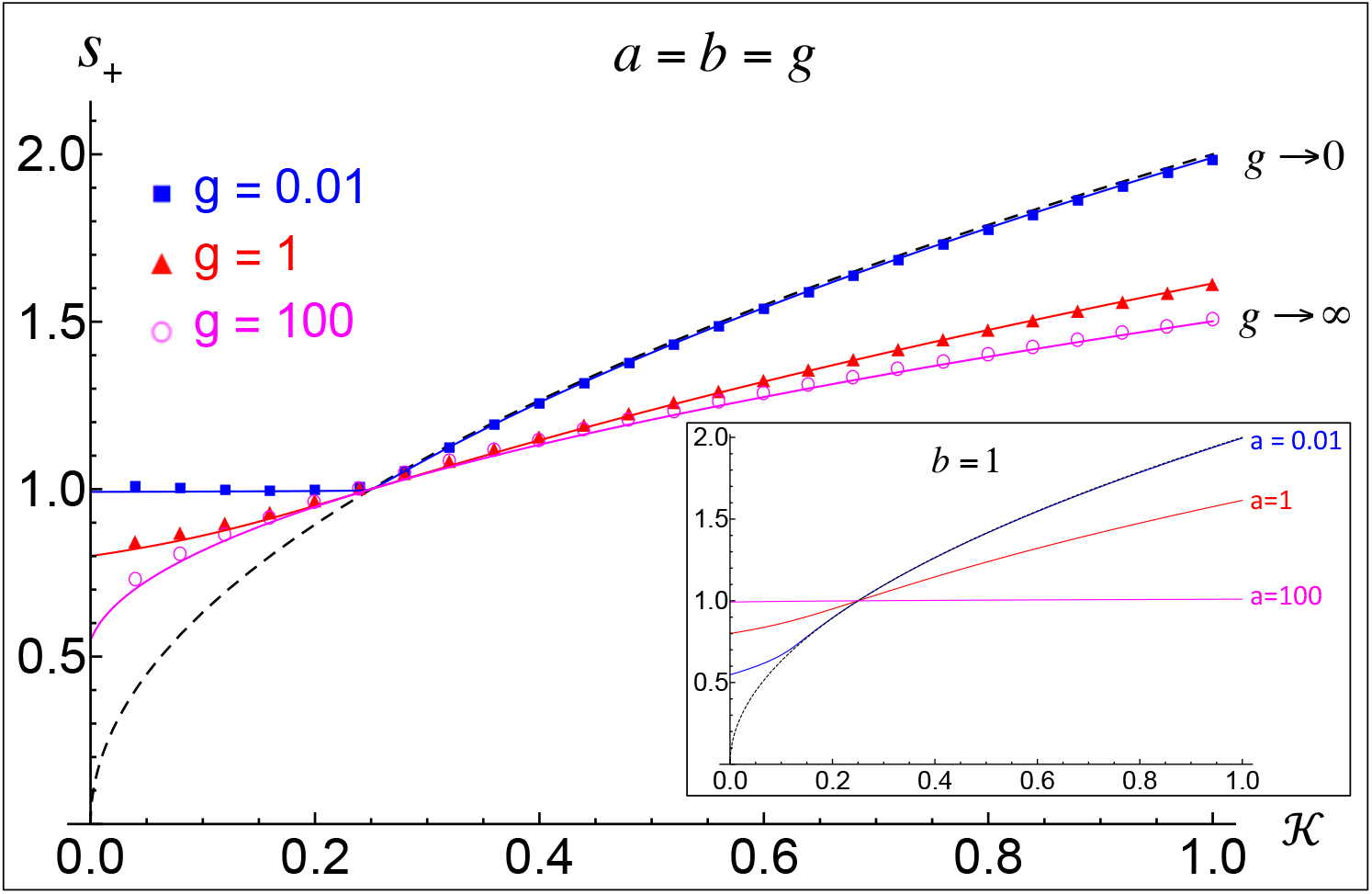
(Color online) The dimensionless speed of the downwind front vs. 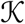 for several coupling values. The symbols were obtained by numerical solutions of Eqs. (3)–(4), while continuous curves are theory – see [33]. Dashed curves are 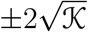 – FKPP speed in units of vo.

On the other hand, there is no drastic change in going from 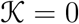, to an infinitesimal 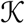. In other words, the limit 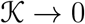 gives the same result as 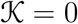 given by Eqs. (7)–(8); adding a diffusive term to an advection-only model, modifies results gradually as 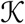 is increased. And for some parameters (such as *a* = *b* ≪ 1) there is essentially no change up until a critical value of 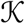.

Both of these observations lead us to conclude that the advective layer can not be ignored. A diffusion-only model infinitesimally perturbed by adding a competing advective transport channel gives rise to a qualitative, finite change in results. On the other hand, advection-only model infinitesimally perturbed by adding a diffusive term in the GL gives rise to an infinitesimal changes in results. Advective layer matters at any coupling!

To stress this point further, we note that at sufficiently-large 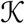, the speed scales with the square root of 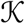 for any *a* and *b*. However, the prefactor does depend on *a* and *b*. Thus, the advective channel affects the results even at arbitrarily large 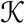.

A very important point is in order. Here we are dealing with dynamics of two variables – *ρ* and *σ*, each describing the densities in advective and diffusive layers respectively. The transport mechanism in the former is advection, and the transport mechanism in the latter is diffusion. In other words, a particle is *either* advecting, *or* diffusing. Had we considered a single layer, with a single density variable, and both mechanisms present in that one single layer, the results would have been completely different. In such advect *and* diffuse scenario (FKPP model with advection), the front speed is simply the advective speed plus the Fisher speed. This can be seen by switching to a co-moving reference frame and thereby removing the advective term from the equation. In our two-layer advect *or* diffuse model, however, one can not get rid of advection by any such transformation. In other words, this model describes a genuine *competition* between the two mechanisms – unlike the FKPP model with advection. The presence of this genuine competition between the two transport mechanisms is what gives rise to all the richness of the model considered here.

## 3 Adding biological realism

The phenomenon of fragility is significant. It suggests that advective transport channel, such as the free atmosphere, can not be ignored – even if the probability of reaching that transport channel is infinitesimal.

This theory, however, is incomplete. We estimate parameters *α, β, δ*, and 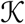 in Appendix A, and conclude that for realistic physical scenarios, *a,b* ≪ 1. Thus, according to Fig. 4, our basic theory predicts the front speed that is at least as big as the wind speed, or even greater if 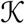 is large enough (see discussion of 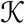 in Appendix A). Traveling at wind speeds typically found in the free atmosphere (see Appendix A), the front would advance by hundreds of kilometers per day. On the other hand, observed invasions of fungal epidemics typically move tens of kilometers per day [1], [26] (see Chapter 14), [45]. Therefore, the simplistic theory in [33] is overestimating the front speed by an order of magnitude.

There are two biological effects that were not included in the model. The first is the role of spore death during the atmospheric transport. For instance, Aylor has shown [1] that the fraction of spores that remain viable after a continuous transport for 700 km can be as low as 10^-12^, although it could be much higher – up to about 10^-2^ if traveling under cloudy conditions. Another factor that may potentially impact viability of spores that travel in the free atmosphere for prolonged periods of time is dehydration [26].

The second piece of missing biology is the fact that spores don’t produce new spores directly. Spores produce fungi, which in turn produce spores [46], [47]. A good, and perhaps the most relevant example for long-range spread is the case of rusts [48]. During the summer invasion event, new fungi reproduce asexually. The asexual cycle consists of infection of a new plant uredospores, followed by a latent period of time, during which the new fungus matures, followed by the onset of production of new uredospores. A typical latent period for rusts is of the order of one week. Thus, a more realistic model would depart from the simplistic treatment of spores as “self-reproducing dust” and would instead take into account both spores and fungi, as well as a delay (latent) time period.

To incorporate both effects, we will now consider the following augmented model. The variable *ρ* represents the density of spores in the AL, as before. The variable *σ* represents the density of spores on the ground and in the ABL, as before. The density of fungus on the ground will be represented by a new variable *φ*. The spores have a probability per unit time *γ* to invade the plant and produce a new fungus after a latent delay period *τ*. The spores also have a probability per unit time Δ to die. All together, the new model will be

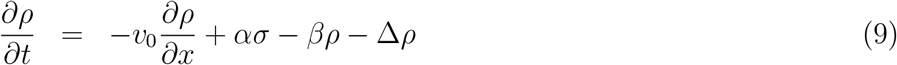

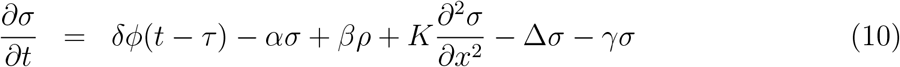

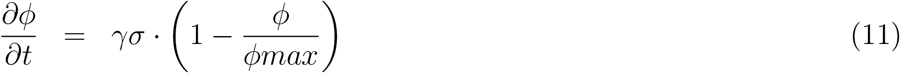

Note that now the growth term is different: only a mature-enough fungus can contribute to new spores. Thus, the addition to the spore density at time t is determined by the fungal density at time *t* — *τ*. Another change is that there is no more carrying capacity for spores, but there is carrying capacity for the fungus, *φ_max_.* This is a more realistic model of carrying capacity than in the one used in [33]; the spores reside on the surface of plants and on the ground, while the fungus resides within the host plant, and its density is subject to carrying capacity set by the host.

To keep the model as compact as possible, we will assign the same death rate in both layers for now. The effect of different spore death rates will be examined in Section 3.2, with the conclusion that the observed difference (no more than a factor of 2, between the ground and 3 km above ground) does not change our conclusions. However, even when the rates are set to the same value, the UV in the AL may still affect spores more than in the GL because the exposure time is greater in the AL. We will examine if this is true also.

Finally, the meaning of *δ* is now different than before. Whereas in the basic model, *δ* is the exponential growth rate, here it is the rate at which the fungus spews out spores (see [26] for the biological details of this fascinating process).

Working again with a constant velocity *v*_0_, and rescaling time by 1/*δ*, and space by *v*_0_/*δ* we have (again, re-using letters *x, t, τ ρ, σ*, and *ϕ* for readability):

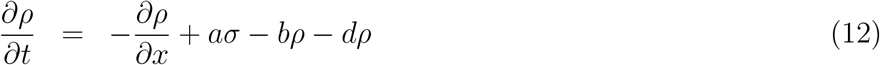

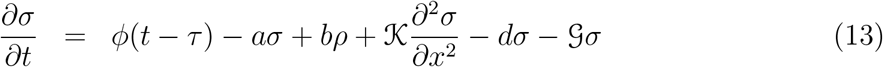

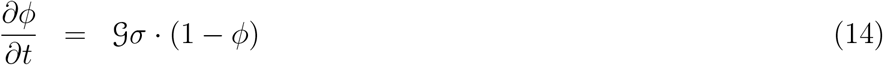

(note the correspondence 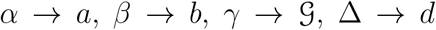 (stands for spore death); the letter *g* has already been reserved for the case of *a = b*). We will see again that fronts in this model are pulled – that is, numerical simulations of these equations will match with analytical calculations in which the term 1 – *ϕ* in the last equation is dropped, making the equations linear.

### 3.1 Does the fragility effect remain?

The key effect underpinning the predictions of the basic model was fragility – a finite change in the front speed due to an infinitesimal coupling to the advective transport channel (AL). If this fragility remains, it implies that AL remains the dominant transport mechanism. If the fragility disappears, it will indicate that AL can be ignored.

For the remainder of the paper, we will focus only on the downwind front speed, so the subscript + will be dropped from *s*_+_.

First, it is easy to find parameters for which fragility certainly does remain. Fig. 5, for instance, compares the front speed vs. 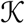 when *a = b* = 0 (diffusion only transport) – red dots, with the case when a competing transport channel has been opened by introduction of a small *a* = *b* = 0.01 – blue dots. The other parameters were *τ* =1, 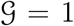, *d* = 0. For this particular parameter combination the two branches merge into one – we see that after 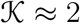 the red and the blue dots lie on the same curve, but such merging is not necessarily the case for all parameter combinations.

**Figure 5:**
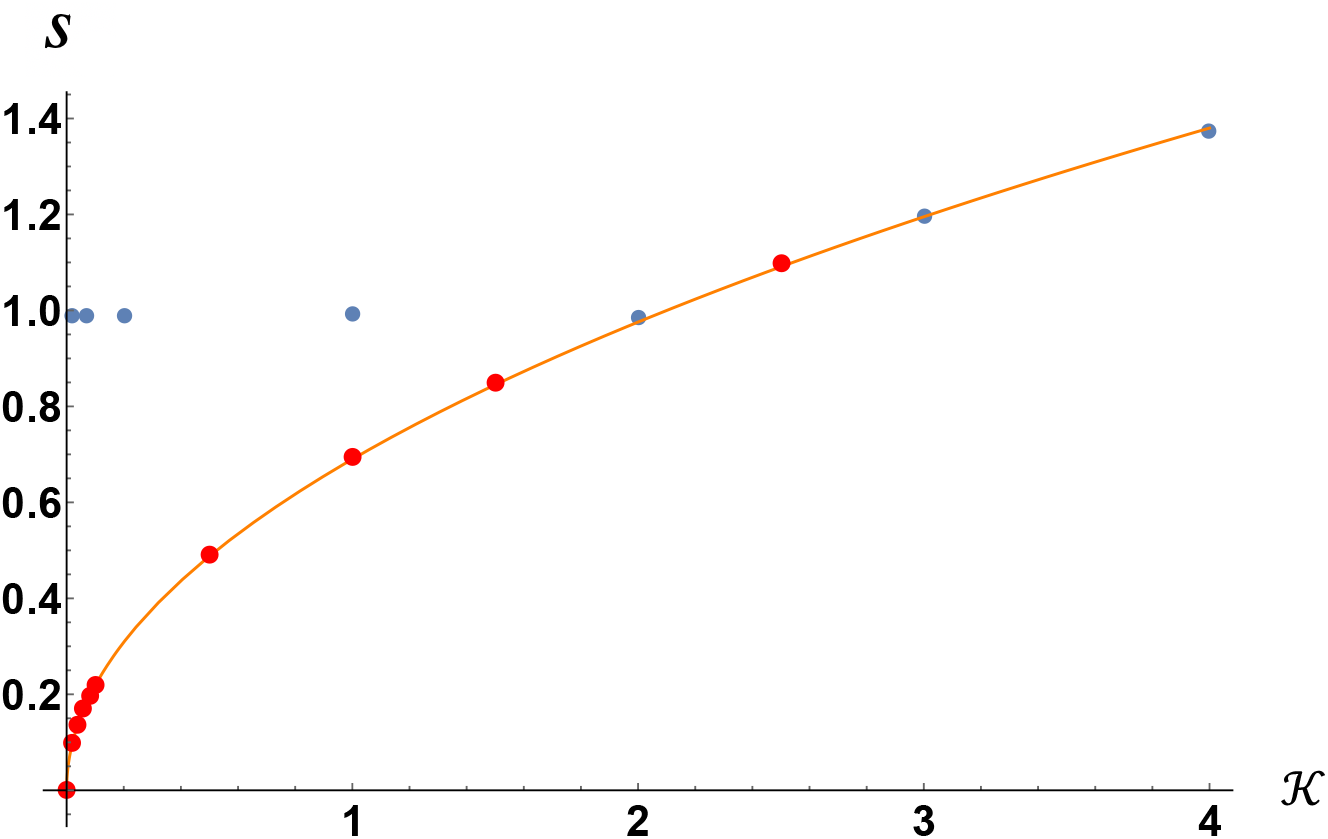
(Color online) Blue dots: *s* vs. 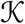 at *a* = *b* = 0.01. Here *τ* =1, 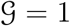, *d* = 0. Red dots: *a* = *b* = 0. Orange curve is a fit to the red dots, given by 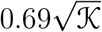. These results are obtained with simulations. See Appendix B.3 for comparison with analytical predictions.

We learn two things from this figure. First, that the square root scaling of the diffusion-only branch remains when latency is present. This is actually just a consequence of dimensional analysis – in the absence of advective mechanism, the front speed has to scale *as* 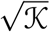 purely on dimensional grounds. Second, we recognize the familiar behavior – c.f. Fig. 4. There is a diffusion-only branch, a coupled branch, and there is a qualitative change in *s* vs. 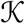 due to the opening up of a competing advective channel by a small *a* and *b*; the system switches from following the diffusion-only branch to following the coupled branch.

To see if the difference between the branches approaches a finite value as *a = b* go to zero, we fix the value of diffusion coefficient at 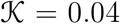, and allow *a = b* to vary. This is depicted in Fig. 6, for several values of *τ*.

**Figure 6:**
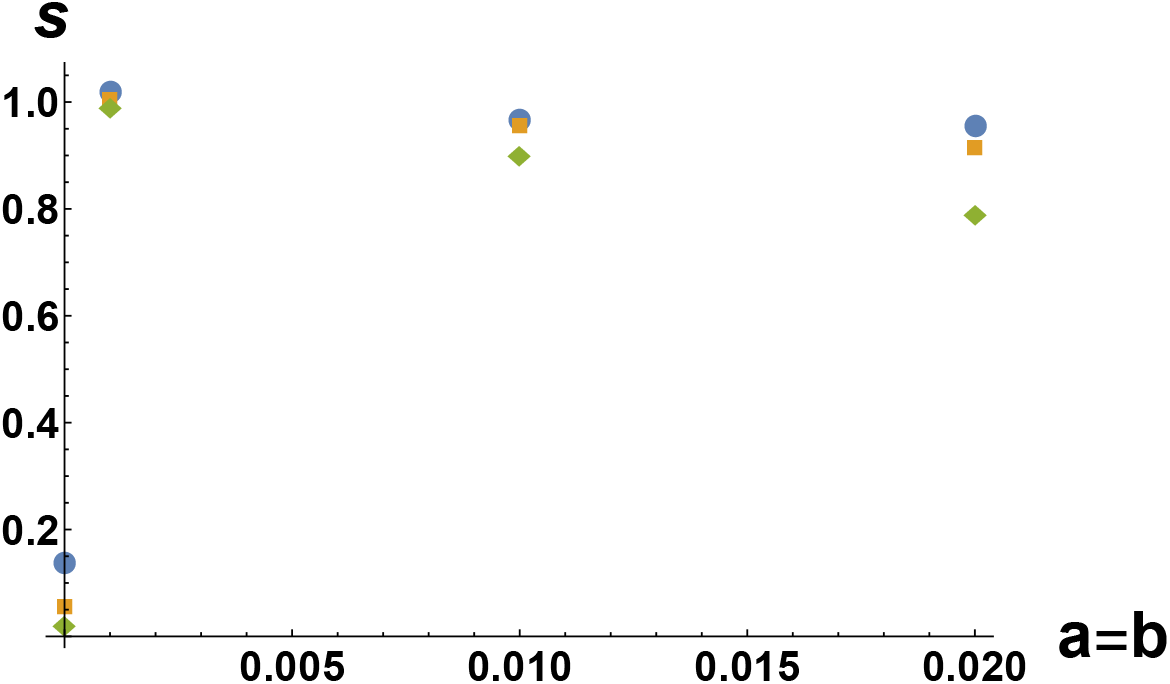
(Color online) *s* vs. *a = b* at 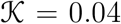, and various latent times. Filled circles: *τ* = 1, squares: *τ* = 10, diamonds: *τ* = 40. Here 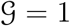, *d* = 0.

The abrupt jump in the speed is evident for all *τ* values considered; extrapolating to *a* = *b* → 0, we get a different number than at *a = b* = 0. This is fragility – turning on infinitesimal coupling to the AL modifies results by a finite amount. The curve tends to the finite value as *a = b* → 0 without any evidence of a decreasing trend. Note that there is no need to look at larger values of 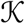; it is at the smallest 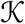 is where fragility is going to be most prominent.

Next, we perform a similar study with a fixed *τ*, but at various spore death rates. This is depicted in Fig. 7.

**Figure 7:**
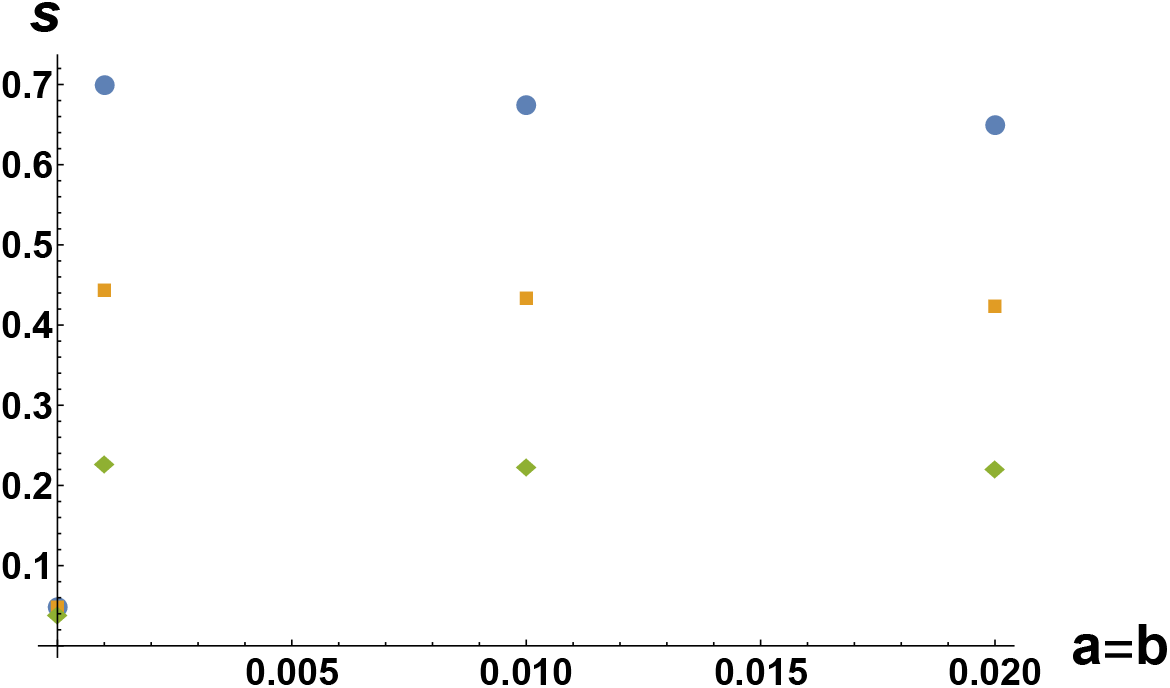
(Color online) *s* vs. *a = b* at 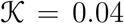, and various spore death rates. Filled circles: *d* = 0.07, squares: *d* = 0.2, diamonds: *d* = 0.5. Here 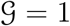, *τ* = 10.

There is still fragility – extrapolating to *a = b* → 0, we get a different number than at *a = b* = 0. However, the discontinuity decreases with increasing spore death rate, and may possibly disappear at even larger *d*. The range of d shown here reflects the estimates described in Appendix A.

Third, we vary 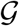. This is shown in Fig. 8.

**Figure 8:**
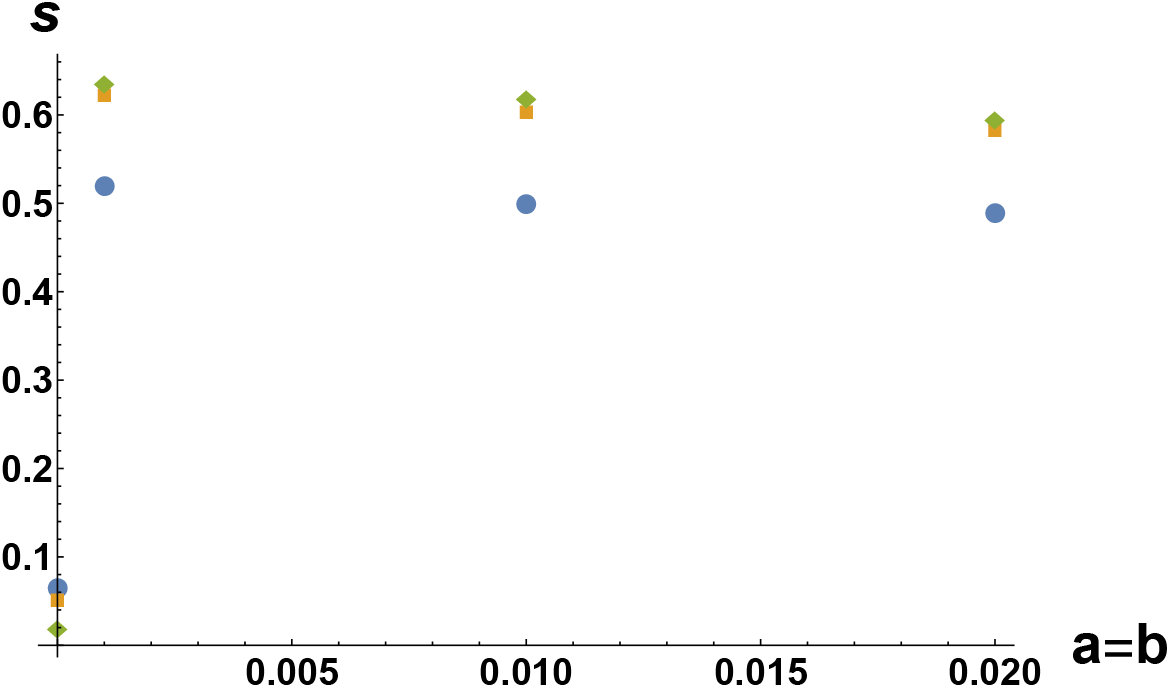
(Color online) *s* vs. *a = b* at 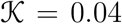, and various values of 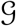. Filled circles: 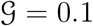, squares: 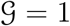, diamonds: 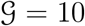. Here *d* = 0.1, *τ* = 10.

This plot suggests that fragility may possibly disappear at a small-enough value of 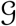. Realistic 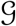 is indeed much less than 1, but *τ* is also much greater than 1. In Appendix A, we estimate that *τ* is on the order of 10^5^ in our time units. Thus, we need to understand the limit of very large *τ* and very small 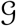. Such large *τ* are completely inaccessible to simulations, due to dramatic increase in simulation time with *τ*. It is in this circumstance where theory becomes invaluable.

Thus, we will now turn to theory for two reasons: first, to explore parts of parameter space that are inaccessible to simulations, and second, to conduct a systematic exploration of parameter space in order to understand whether fragility remains for all parameter combinations. We verified the theory on those parameters where theory and simulation could be compared – see Appendix B.1 – B.2.

As parameters change, both the coupled and diffusion-only branches will evolve. While the value of the coupled branch at a given 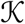 could decrease, this does not indicate that the advective mechanism is becoming less important because the value of the diffusion-only branch at the same 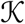 could also decrease. Therefore, we must compare the diffusion-only branch – one that gives the front speed at *a = b* = 0, with the coupled branch – one that gives the front speed at a small, but non-zero value of *a = b*. In the results that we now present, we chose *a = b* = 10^-3^. The value of 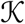 has to be held fixed as the coupling is turned on or off. We will present results for a range of 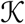.

We varied 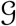 (rate at which spores infect host plants), for a set of different values of spore death *d*, while *τ* was fixed at 10^4^. We will comment on the effect of changing *τ* below. At each combination of parameters, we plot the value of both the coupled, and the diffusion-only branches at that 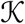. These results are presented in Fig. 9. The curves in these figures come from the solutions of the transcendental Eqs. (39)–(40). The details of this procedure can be found in Appendix B.3.

**Figure 9:**
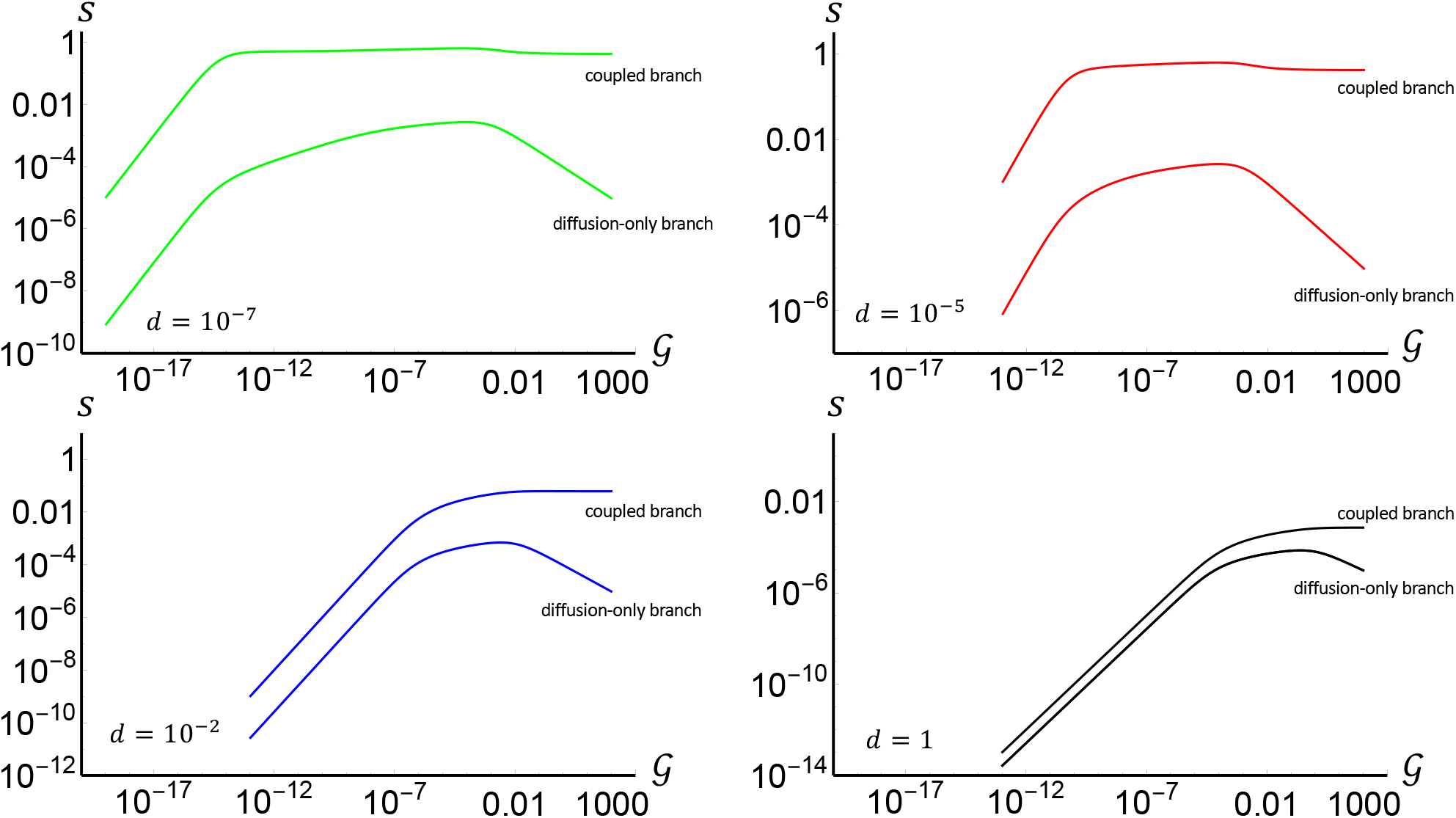
(Color online) Coupled and diffusion-only branches for a range of infection rates 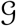, at various spore death rates *d*. The coupled branch corresponds to the coupling *a = b* = 10^-3^ between the AL and the GL; the diffusion-only branch is uncoupled from the AL, i.e. *a = b* = 0 along that branch. The diffusion coefficient is 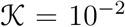 and *τ* = 10^5^. Note, at d =1, the speed of the coupled branch is at least 3.9 times larger than the speed of the diffusion-only branch.

It is clear that the speed of the diffusion-only branch is smaller – and typically many orders of magnitude smaller – than the speed of the coupled branch at the same 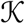. The gap decreases with increasing *d*, but even at *d* = 1 the speed of the coupled branch is at least 3.9 times larger than the speed of the diffusion-only branch. We estimate in Appendix A.1 that realistic values of d will not likely be much above 10^-2^, and for many species of pathogens they can be more than an order of magnitude below even under sunny conditions, and several orders of magnitude lower under cloudy conditions.

In Fig. 9 we used 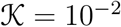. The next figure demonstrates the changes that occur when 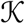 is increased by a factor of 10.

We learn two things. First, we learn that changing 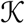 from 10^-2^ to 10^-1^ only affects the diffusion-only branch, and not the coupled branch. We found this to hold true even as 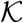 was increased furhter. This information will be used in the next section.

Second, while the narrow gap at *d* = 1 closes, most of the other gaps are changed by a relatively small fraction. We estimated in Appendix A.1 that realistic values of d are below 10^-2^. In order to close the gap at *d* = 10^-2^, 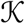 needs to be about 10. To close the gap at even smaller *d* requires much larger 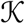. Recall that dimensionless 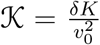. We estimate in Appendix A that *δ* ≈ 400 *hr*^-1^ and *K* ≈ 0.05 *km*^2^/*hr*. In order for 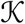 to be of order 0.01, *v*_0_ has to be of order 40 *km/hr*, and in order for 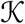 to be of order 10, *v*_0_ has to be of order 1.5 *km/hr*. Recall that *v*_0_ represents the difference between the mean winds in the ABL and the advective velocity in the free atmosphere. A difference of 1.5 km/hr is certainly plausible. This estimate represents extremes. Thus, *d* = 10^-2^ represents an upper range of typical spore death rates, and *v*_0_ larger than 1.5 km/hr is also realistic. Therefore, the gap will remain non-zero for a large range of pathogens and physical conditions.

In the rest of this section we go back to 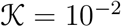. In the next graph, we compare results of *τ* = 10^4^ (solid curves) with *τ* = 10^5^ (dashed curves) at two values of *d*. Same values of *a = b* and 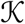 were used as in Fig. 9.

It is also interesting to see which features of graphs are affected by spore death directly. Fig. 12 addresses this. Evidently, at low spore death rates, only the low-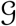 tail is affected by spore death. However, as the spore death rate increases, the entire function 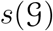 is modified.

**Figure 10:**
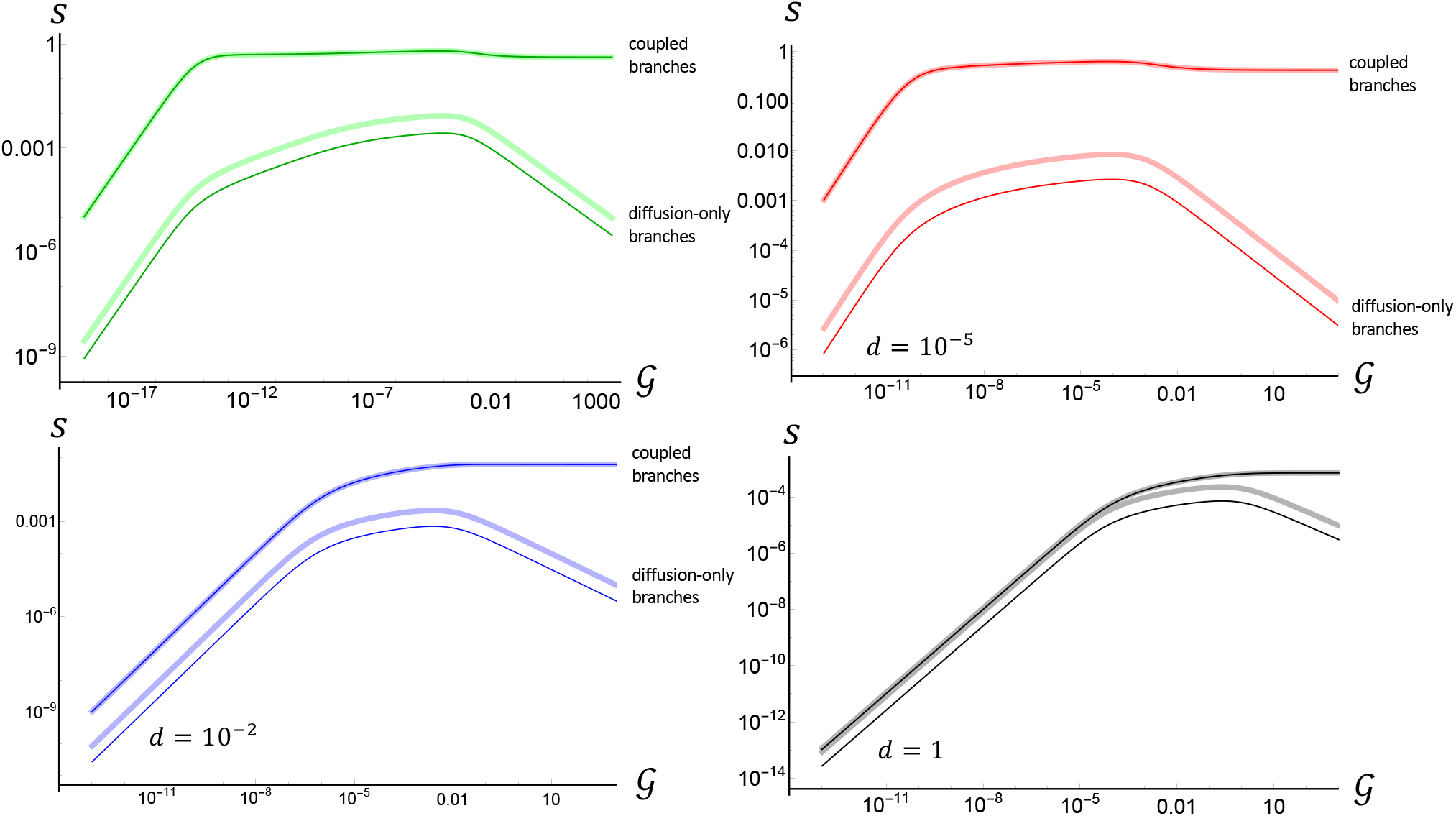
(Color online) Coupled and diffusion-only branches for a range of infection rates 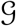, at various spore death rates *d*. Comparison is made between results for 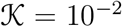 (thin, dark curves), and 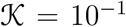 (thick, light curves). The coupled branch corresponds to the coupling *a = b* = 10^-3^ between the AL and the GL; the diffusion-only branch is uncoupled from the AL, i.e. *a = b* = 0 along that branch. **axes**

**Figure 11:**
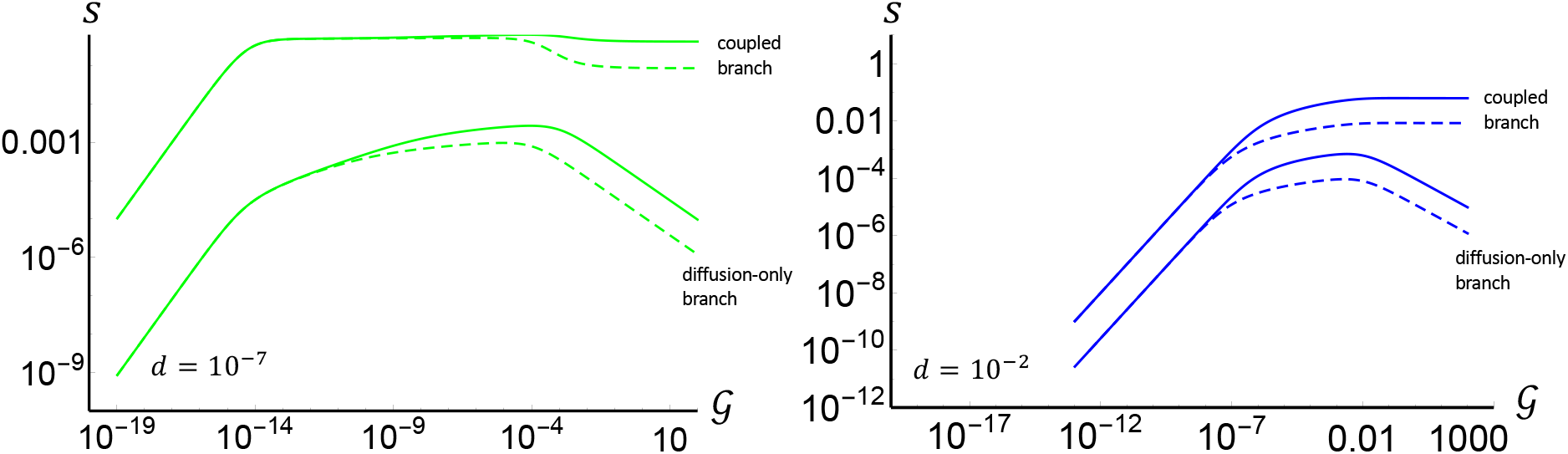
(Color online) Comparison of 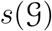 for latent time *τ* = 10^4^ (solid curves) and *τ* = 10^5^ (dashed curves), at two values of spore death rate *d*. Same values of *a = b* and 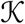 were used as in Fig. 9.

**Figure 12:**
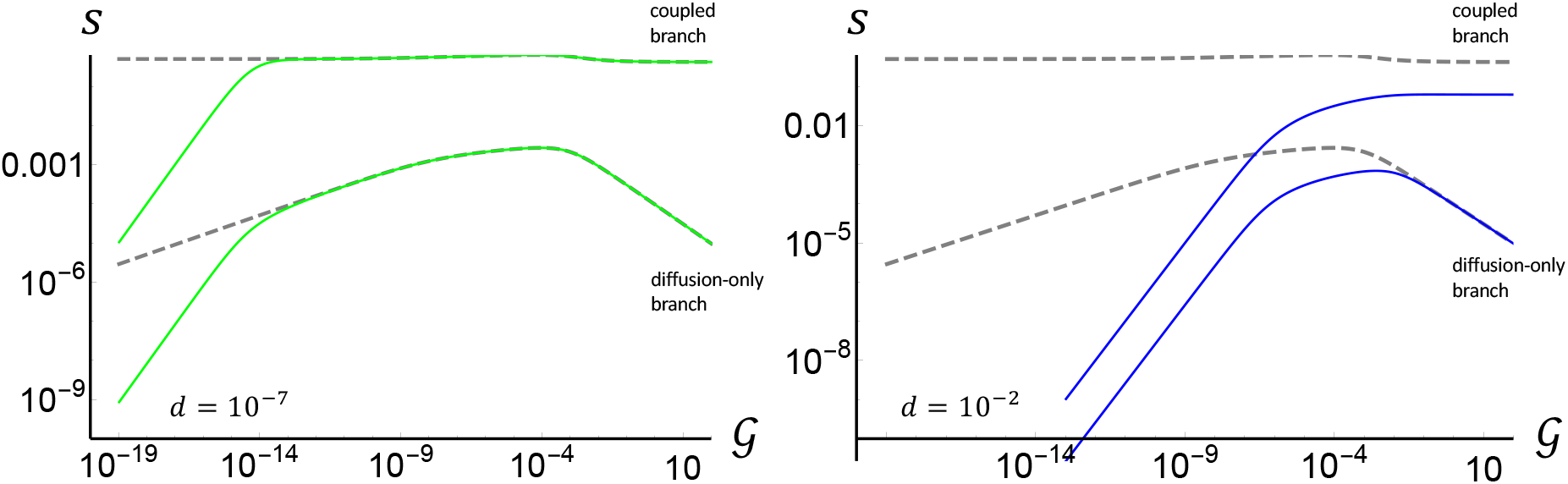
(Color online) Both branches of 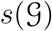 for *d* = 0 (dashed curves); *d* = 10^-7^ and *d* = 10^-2^ (solid curves in left and right panels respectively). Here *τ* = 10^4^. Same values of *a = b* and 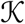 were used as in Fig. 9.

We present one more study.

Fig. 13 compares the gap between the two branches at small and large couplings rates, and for a zero and a non-zero spore death rates. We chose to compare *d* = 1 with *d* = 0, because *d* = 1 is a relatively large spore death rate in terms of what’s actually observed. Also, we chose couplings three orders of magnitude below and above 1.

**Figure 13:**
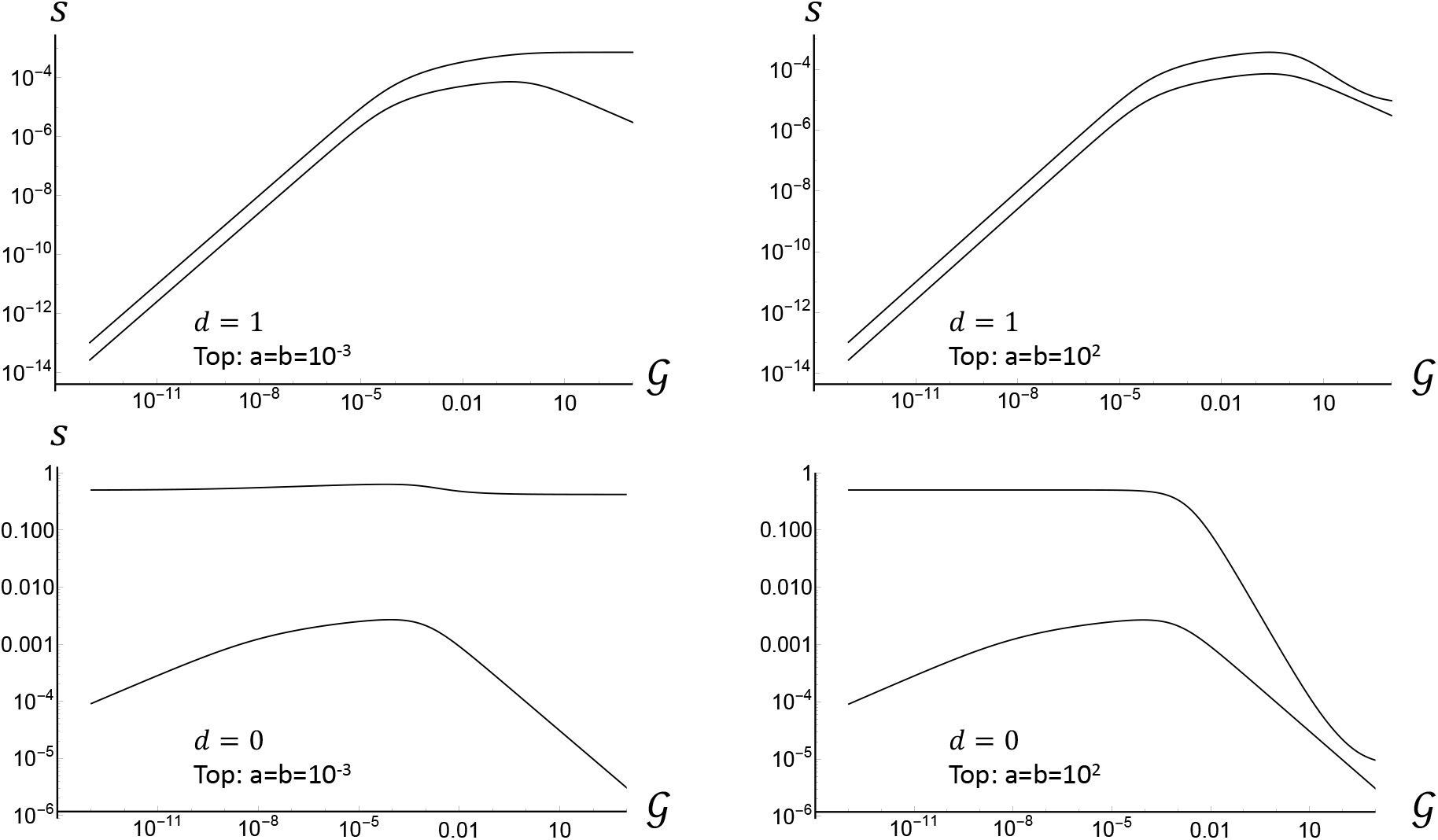
Comparison of both branches at small (left) and large (right) couplings. Top: large spore death rate (*d* = 1), bottom: no spore death (*d* = 0). In all cases, *τ* = 10^4^ and 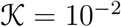.

In going from *d* = 1 to *d* = 0, the values of both branches is increased – fronts propagate faster in the absence of spore death. However, the change that results from the absence of spore death is more dramatic for the coupled branch than for the diffusive-only branch. Furthermore, the coupled branch experiences a change for all values of 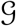 at small couplings, but only for sufficiently small 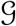 at large couplings. Both of these observations are consistent with our intuition that AL is affected by the spore death more than the GL.

It is worth noting that in going from small couplings to large couplings, there is a widening of the gap at larger 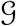. However, this is not related to spore death, since this effect takes place even at zero *d*.

### 3.2 Different spore death rates in the two layers

We now investigate the role of different rates of spore death in the two layers. The intensity of UVB increases with height above the ground. The UVB irradiance is a function of many parameters – including the angle of the Sun above the horizon, atmospheric turbidity, cloud cover, and elevation. The direct component of solar radiation increases with height, while the scattered component decreases. Different sources report different height dependence of intensity in the UVB range. Aylor [26] states that “On clear days, the intensity of global UV is about 30 % higher at an attitude of 3, 000 *m* than at sea level”. On the other hand, Maddison and Manners [54] report that “... the net increase in the 290 — 360 *nm* waveband might be of the order of 15 — 30 % per *km* increase in height” (evidently, both sources refer to the net effect of UVA and UVB). Thus, Maddison and Manners predicts UV intensity at the height of 3 km to be between 1.5 and 2.2× the value at the ground, rather than 1.3× reported by Aylor.

We would like to study the worst-case scenario for the effect of UV. Thus, we will set *d_AL_* = 2*d_GL_*, and for *d_GL_* itself, we will work with 10^-2^, which is also in the upper range of possible spore death rates (see discussion above and in Appendix A.1).

The solid black curves in Fig. 14 are the two usual branches – the coupled branch and the diffusion-only branch, for *d* = 10^-2^. We now compare the two branches with the situation in which the GL spores die at the same rate, i.e. *d_GL_* = 10^-2^, while the AL spores die at a higher rate. The top dashed pink curve corresponds to the coupled branch for which *d_AL_* = 2*d_GL_*. The bottom dashed pink curve corresponds to *d_AL_* = 30*d_GL_*. Although this is far beyond the real *d_AL_*, we used this number to demonstrate how much larger *d_AL_* needs to be in comparison to *d_GL_* in order to close the gap between the two branches. Both dashed pink curves are coupled branches; the diffusion-only branch (*a = b* = 0) stay the same as the diffusion-only branch with *d* = 10^-2^, because the spore death rate on the GL is still 10^-2^. These results are based on solutions to transcendental Eqs. (39)–(40) in Appendix B.4.

**Figure 14:**
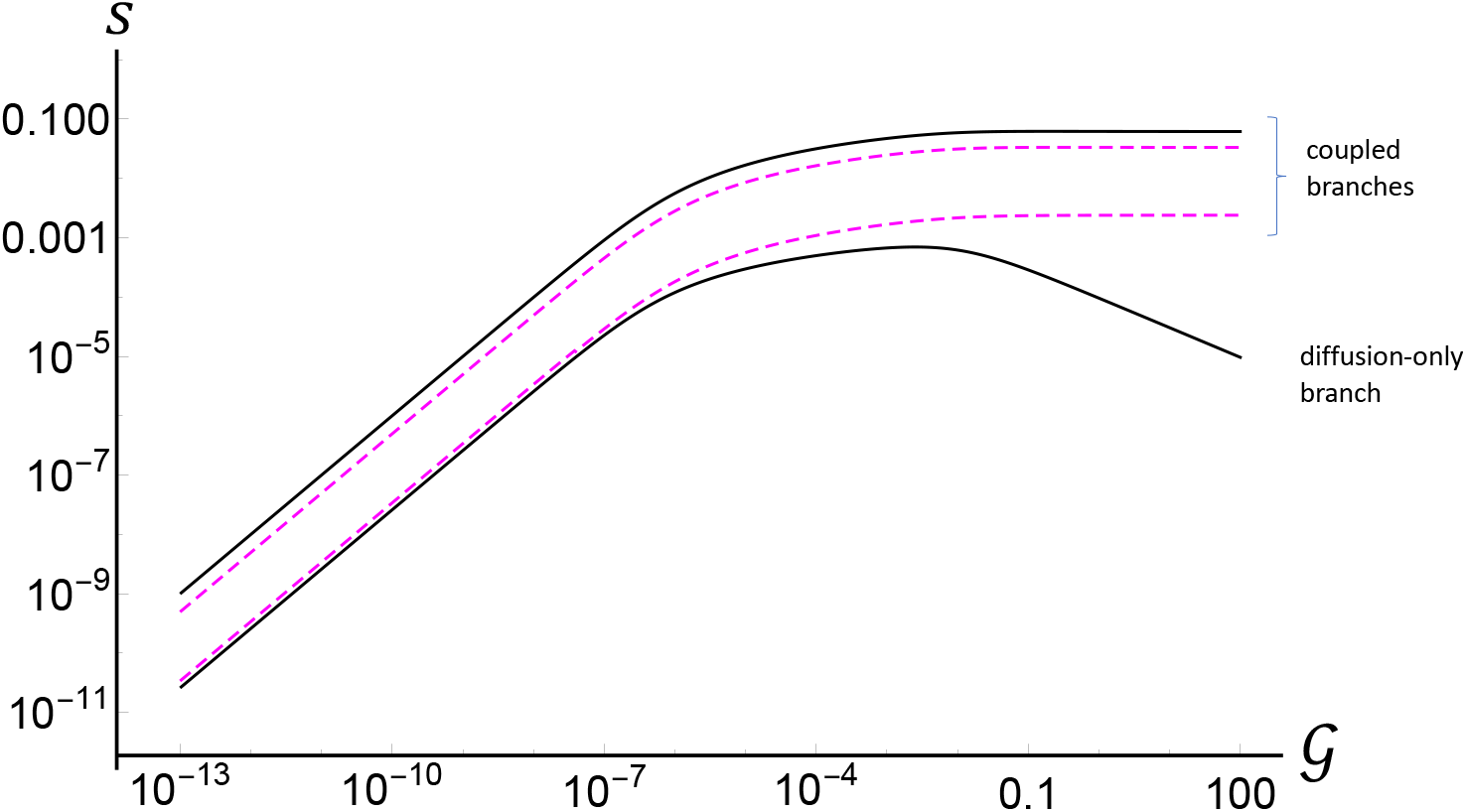
(Color online) Solid black curves: *d* = 10^-2^ in both layers. Dashed pink curves: coupled branches with *d_AL_* = 2*d_GL_* (top dashed curve) and *d_AL_* = 30*d_GL_* (bottom dashed curve); *d_GL_* = 10^-2^ in both cases. Here *a = b* = 10^-3^ on coupled branches, 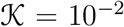, and *τ* = 10^4^.

For *d_AL_* = 2*d_GL_*, the gap between the coupled branch and the diffusion-only branch has decreased by a small fraction of its value with identical spore death rates. The gap shrinks as *d_AL_* increases. In order for the gap to decrease by a significant percentage – say 80% of its value at identical spore death rates, *d_AL_/d_GL_* must be much greater than 2 – an upper limit of a value that characterizes a realistic physical situation.

What happens in the limit of infinitely large *d_AL_*? To check this, we took an analytic limit *d_AL_* → ∞ in Eqs. (39)–(40). The resulting Eqs. (41)–(42) have only one solution branch, which generally lies below the diffusive-only branch – unless *a* = 0, in which case it matches the diffusion-only branch of the model with identical spore death rates. This is expected – removing spores from the GL is equivalent to increasing the spore death rate, since these spores are immediately killed once they arrive to the AL, and will not return back to the GL. In fact, note that in Eqs. (41)–(42), *d_GL_* and *a* appear additively.

We now consider an unrealistically large spore death rate on the GL. The black, solid curves in Fig. 15 are the two branches for *d* = 1. We again compare the two branches with the situation in which *d_GL_* is the same, while the AL spores die at a higher rate. The dashed pink curve corresponds to the coupled branch for which *d_AL_* = 2*d_GL_*. The diffusion-only branch is the same as the diffusion-only branch with *d* = 1, because the spore death rate on the GL is still 1.

**Figure 15:**
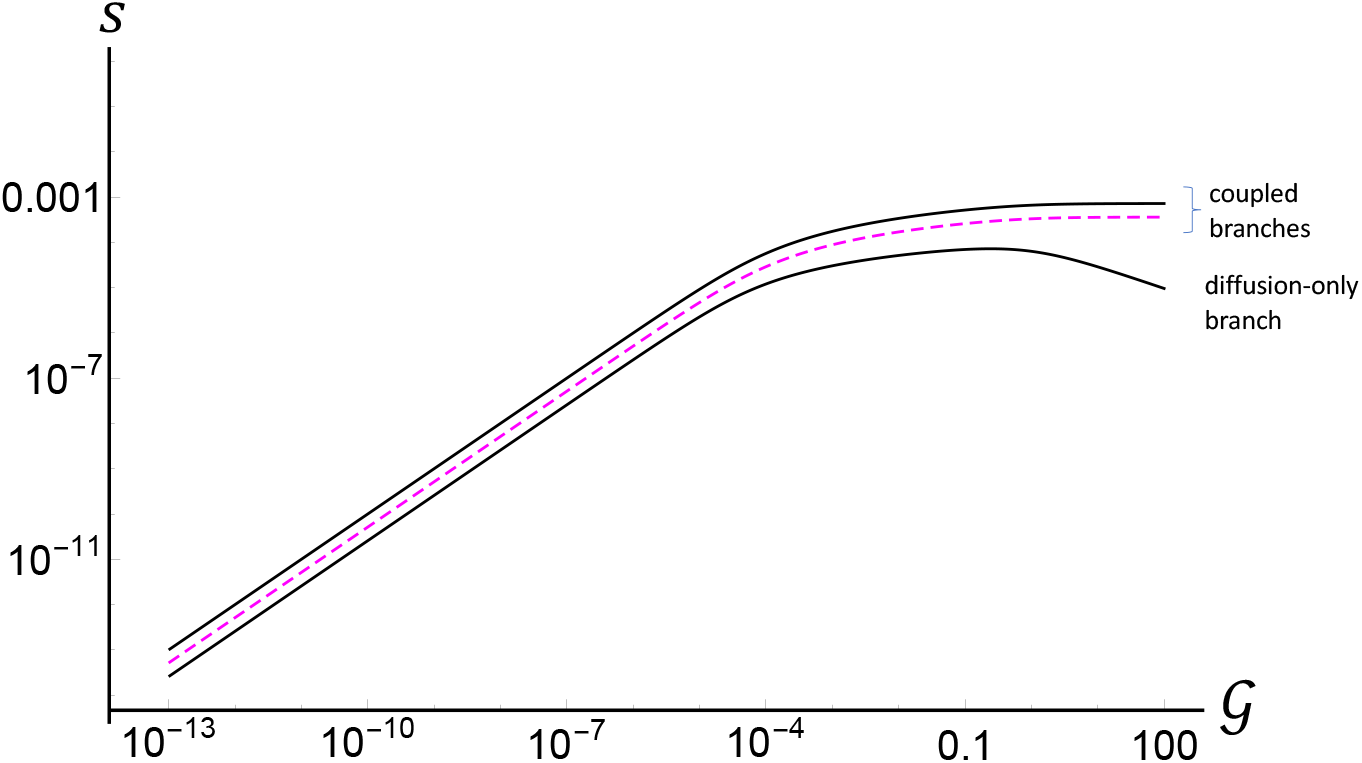
(Color online) Black, solid curves: *d* = 1 in both layers. Dashed pink curve: coupled branche for *d_AL_* = 2*d_GL_* (again, *d_GL_* = 1). Here *a = b* = 10^-3^ on coupled branches, 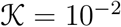, and *τ* = 10^4^.

Based on these results we can conclude that while it is possible to erase the gap between the coupled branch and the advective-only branch with a large-enough *d_AL_/d_GL_*, the typical value of this ratio (below 2) is such that a substantial gap remains for realistic values of *d_GL_*. In short, advective layer matters despite the observed differences in the UVB intensity between the ground and several kilometers above sea level.

### 3.3 Does the new model predict a lower front speed?

One of the problems with the basic model was over-prediction of the front speed. The question is, whether including the (i) spore death and (ii) latency would be able to predict the reduction of the front speed. The answer is yes. We will now support this, and explore how these two factors contribute to the reduction in the front speed.

In Fig. 16 we present a systematic study of how the front speed depends on the infectivity rate 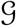, spore death rate *d*, and latent times *τ*. These results are values of the coupled branch. At such parameters, the basic model predicts the speed of the front very close to 1, i.e. essentially at the speed of the wind in the advective layer. So, it is evident that the new model is capable of predicting slower speeds.

**Figure 16:**
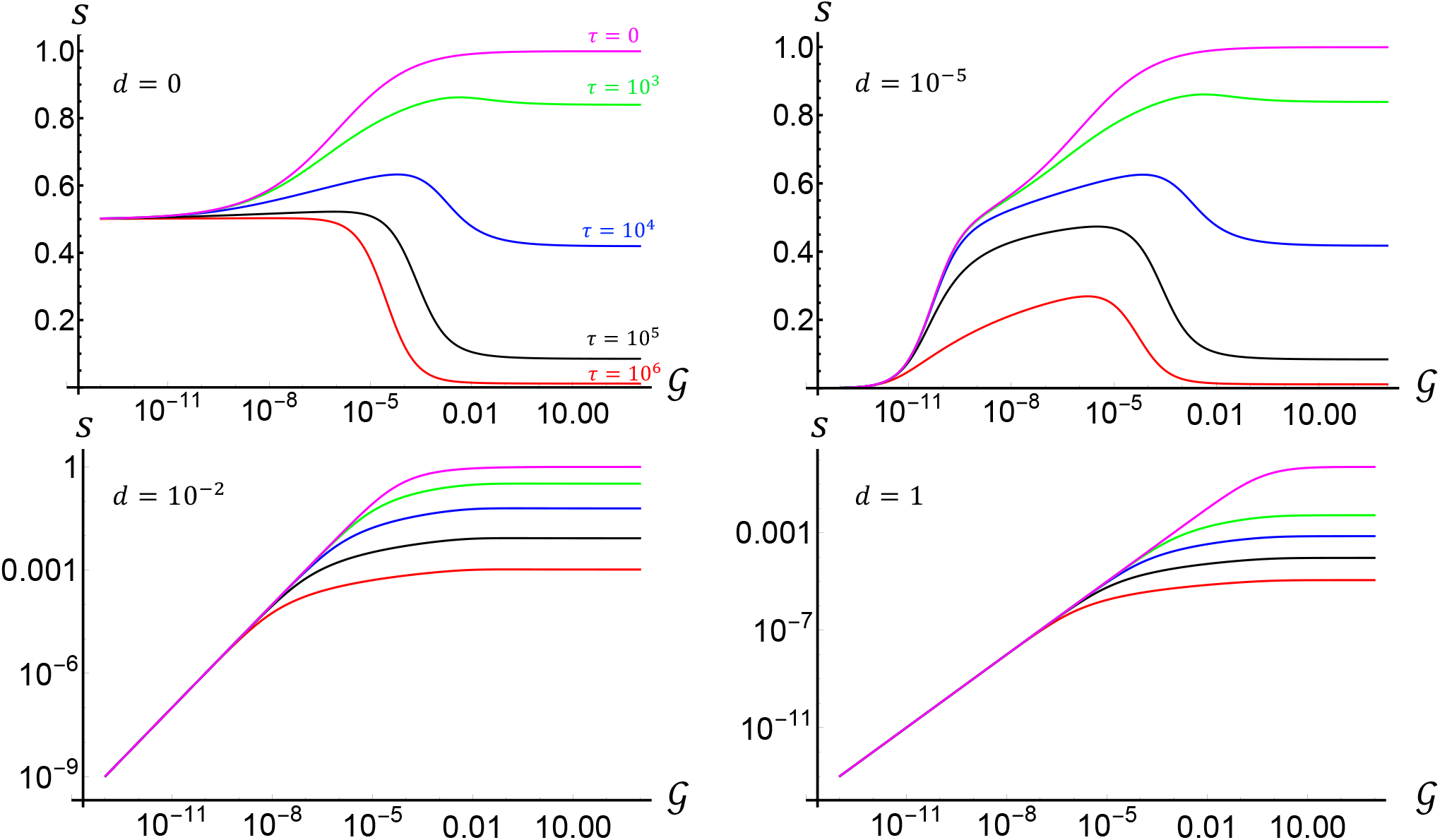
(Color online) Front speed vs. 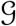 for various spore death rates *d* and latent times *τ*. Here the coupling rates are *a = b* = 10^-3^ and diffusion coefficient is 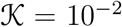. In all of the four graphs, the five curves correspond to *τ* = 0, *τ* = 10^3^, *τ* = 10^4^, *τ* = 10^5^, and *τ* = 10^6^ from top to bottom. Note the linear scaling of the vertical axes in the two upper panels, and logarithmic scaling in the two lower panels.

Note the speed approaching 1 at zero spore death, as *τ* tends to zero and at large 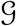. It is a limit in which the new model is most similar to the basic model (although not the same, because the spore production terms are different – see [49]), and the result is indeed consistent with the basic model. On the other hand, at zero spore death and at small 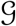 (and any *τ*), the speed approaches 1/2. Zero 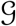 means the spores are not reproducing, and the process reduces to a spread of a pulse of spores with the average of the speed on the GL and the AL, i.e. 1/2. In fact, the basic model with no growth term makes the same prediction [33].

We noted in the previous subsection, that increasing *d* from 0 first affects the low-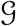 tail of 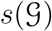, but at larger *d* this function is modified completely from the *d* = 0 form. We can see from the upper-right panel of Fig. 16 that at at *d* = 10^-5^, this low-end tail with value 1/2 is suppressed, but the rest of the function remains intact. The fact that sufficiently low spore death only affects the low 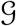 portion of 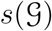 allows us to separate the role that the spore death and latency play in slowing down the front. The two upper panels of Fig. 16 – at *d* = 0 and *d* = 10^-5^ clearly show that increasing the latency decreases the speed. This is expected. The two lower panels include the effect of both parameters, but it is still always true that increasing the latency decreases the speed. This effect is intuitive.

We also slice the parameter space by holding the *τ* fixed at 10^5^ and varying the coupling rates. This is shown in Fig. 17. These graphs demonstrate a non-intuitive feature – that increasing the coupling rate actually decreases the front speed. This effect appears to hold true for any spore death rate (see also Appendix B.2, Fig. 23, corresponding to zero diffusion and zero spore death rate). The same effect was also present in the basic model in the absence of diffusion – see the left panel of Fig. 3. Recall that times are rescaled by growth rate in both models. Therefore, smaller dimensionless coupling rate can be achieved by either decreasing the physical coupling rate, or in equal measure by increasing the growth rate. It is intuitive that a larger growth rate would give rise to a faster front. On the other hand, increasing the physical exchange rate (coupling) while holding growth rate fixed allows less chance for spores to reproduce, so this also lowers the front speed.

**Figure 17:**
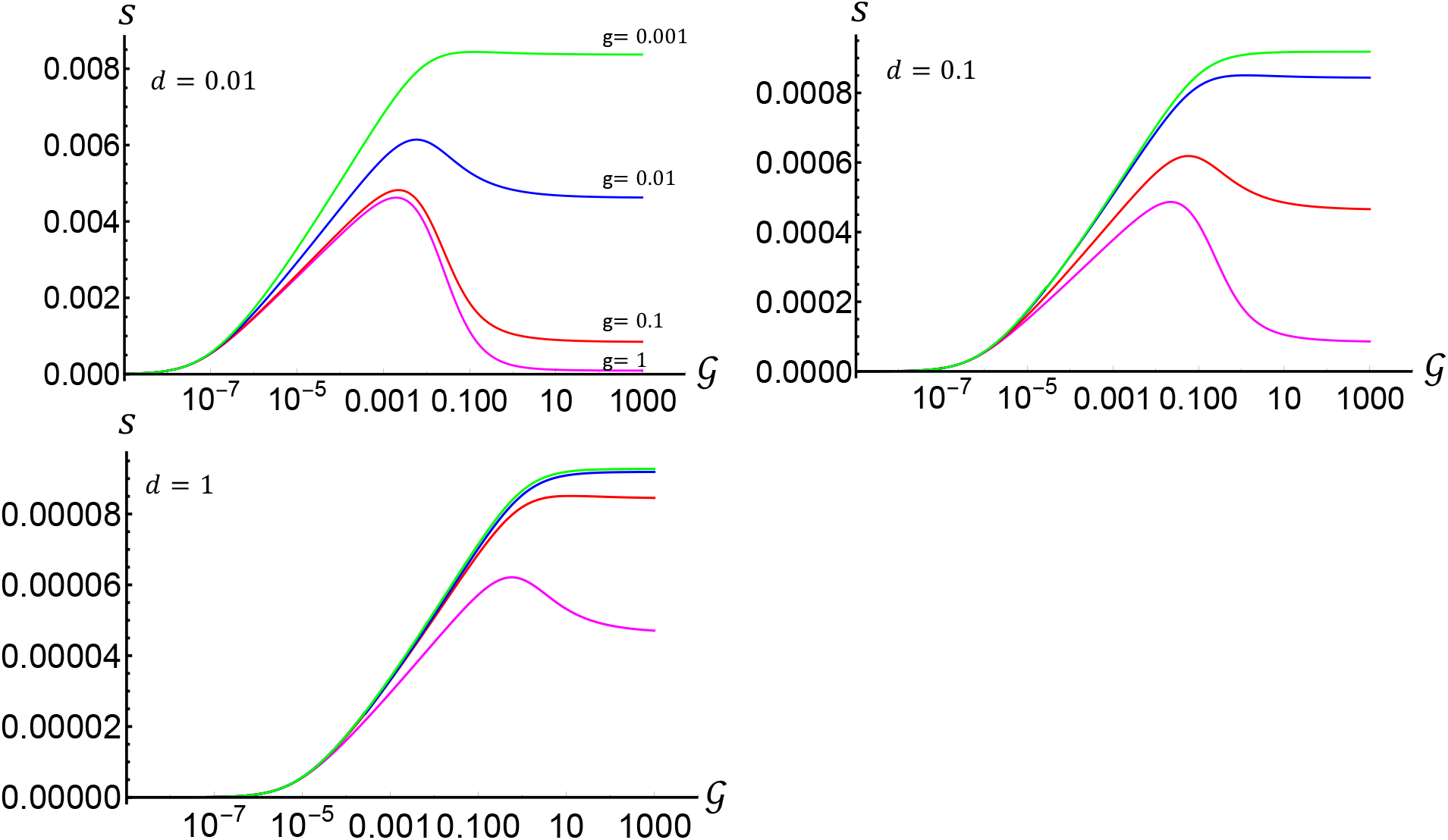
(Color online) Front speed vs 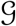 for various spore death rates *d* and coupling parameters *a = b = g*. Here the latent time is *τ* = 10^5^ and diffusion coefficient is 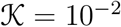.

We estimate parameters in Appendix A. We find that an upper range for *d* is 0.02, 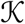 is likely to be between 0.01 and 0.1, and *τ* is on the order of 10^5^. The coupling values *a* and *b* are likely to be much less than 1. With *d* = 0.01 and *a = b* = 0.001, our calculations suggests an upper limit on the speed to be around 0.01, i.e. 0.01 of the speed of the advective layer. Decreasing d by a factor of 10 increases the upper limit to ≈ 0.05 of the advective layer speed. On the other hand, it is possible to have no reduction in the speed – this happens for d below 10^-4^ and for a range of 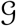. We conclude that for a wide range of parameters there will be a reduction of speed, but to make a more accurate prediction we need a more precise knowledge of parameters.

## 4 Conclusion

In examining continental-scale plant pathogen spread, it is often assumed that only the atmospheric boundary layer (ABL) plays an active role in dispersing spores. This is a reasonable belief, due to the low probability for spores to reach higher layers of the atmosphere and the deleterious effect of UV radiation on spores. In this paper, we examined this belief quantitatively, using a model that explicitly includes the two mechanisms – short-range hopping provided by the ABL and long-range advection provided by the higher troposphere. This model also includes the spore death due to UV exposure. We find that in the competition between the inefficient but safe short-range hopping mechanism, with the efficient but improbable long-range advective mechanism, the free troposphere would, in fact, be the crucial mechanism. That is, even if the probability of spores reaching it and surviving within is infinitesimally small, it remains a mechanism that can not be ignored. Ignoring it would lead to results that change by a finite amount even if the coupling to this layer is infinitesimal. We called this fragility. The quantitative measure of fragility was a characteristic size of the gap between the coupled branch and the diffusion-only branch in the function of front speed versus the diffusion coefficient 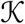. While the gap can be closed in some regions of parameter space, it remains substantial in the realistic parameter range. Even when we looked at worst case scenarios in the far reaches of realistic parameter values, the gap still remains substantial.

Thus, our results challenge the commonly held belief, and we claim that the upper atmosphere can not be ignored – despite a small chance for spores to reach it, and despite a longer exposure times to UV. The short-range hopping mechanism that is typically considered in phytopathology [18] – [23] is fragile with respect to perturbation by an alternative advective mechanism.

## A Estimation of model parameters

Before estimating dimensionless parameters, we need to relate the physical units of time in seconds to the dimensionless units used in this paper.

Here we work with dimensionless time *t*′ = *δt*, where *t* is in seconds, and *δ* is the rate of spore production per uredium postule per second. Using Wheat Stem Rust as a reference, one uredium postule produces on the order of 10^4^ uredospores per day – see Ch. 1 of [48] and references therein. Before normalizing by the carrying capacity, the dimensional density variable *ϕ* as it is used in Eq. (10) represents the number of uredium postules per area, so *δϕ* gives the rate of uredospores produced per area, per day. We note that *δ* of 10^4^ spores per uredium per day is equivalent to ≈ 0.1 spores per uredium per second. Therefore, one unit of dimensionless time, i.e. Δ*t*’ = 1 corresponds to 1/*δ* ≈ 10 seconds, in the case of Wheat Stem Rust.

We can now discuss the latent time in dimensionless units. A typical latent time for Wheat Stem Rust is between one and two weeks, Ch. 1 of [48]. Thus, in our dimensionless units, *τ* is on the order of 0.5 × 10^5^ to 1 × 10^5^, to one significant digit.

Next, we estimate the dimensionless diffusion coefficient, 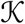. Changing back to physical units, we have 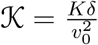, where *K* is a physical diffusion coefficient and *v*_0_ is the speed of the advective layer. The physical diffusion coefficient refers to eddy diffusion [50], not molecular diffusion. Therefore, the value of *K* depends on the type of turbulence relevant to spore dispersal. Empirical data suggests that most spores return back to the ground within 100 meters – see Ch. 13 of [48] and references therein. This distance is much less than a few kilometer scale of the ABL, but comparable to characteristic length of the surface layer in geophysical flow [13] (Chapter 8), where the surface directly influences the properties of the eddies. The turbulence in the surface layer is a wall turbulence, with its characteristic logarithmic mean wind velocity profile in buoyantly neutral conditions [13], [50], [51].

Therefore, we will estimate *K* under the assumption that eddies of this type of turbulence are responsible for eddy diffusion. This approach is similar to other analyses of turbulent-driven dispersal at the plot scale [18], [19] (in this second reference, the authors use a power-law model of velocity profile even for a neutrally-stratified surface layer). However, it does undercount those rare spores that are transported by bigger eddies of the scale up to the width of the ABL itself – same eddies that transport spores to the boundary with the FA and facilitate movement of spores between the layers of our model.

The turbulent diffusion coefficient for near-wall shear-driven turbulence grows linearly with height as *K* = *κv*z*, where *z* is height above the ground [18], [19]. Here *κ* is a universal von Kármán constant ≈ 0.4, *v*\* is the friction velocity ≈ 0.4 m/s [18], [19]. Our model has a constant *K*. We will estimate it as *K* = *κv*L*, where *L* is the width of the surface layer, taken to be 100 meters. When these numbers are substituted, we estimate that *K* ~ 0.05 *km*^2^/*hr*.

To calculate 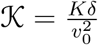 we need an estimate of *v*_0_. In his discussion of transport of a cloud *v*_0_ or plume of spores, Aylor cites “velocity of transport layer” to be 20 — 40 *km/hr* (or 5 — 10 *m/s*) [1], [26] (Chapter 14). Similar figures characterize mean velocity within the mixed layer of the ABL near San Juan, Puerto Rico [52]. Substituting 20 — 40 *km/hr* for *v*_0_, and *δ* = 400 *hr*^-1^, we come up with a dimensionless 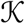 that lies in the range 0.01 — 0.06. Because our method of estimation assumed that only surface layer eddies are responsible for dispersion of spores, we may have under-estimated 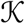 due to rare dispersion events facilitated by bigger eddies. Moreover, the velocity *v*_0_ in our model is not the advective velocity relative to the ground, as in Aylor’s models, but a difference *v_AL_* — *v_GL_*. For this reason, we present results in the main text for a range of 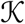. While some graphs will assume 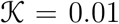, the effect of increasing of 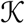 is stated. Most importantly, the 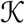 necessary for closing the gap between the coupled branch and diffusion-only branch is also stated.

The parameters *a, b*, and 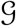 are rather difficult to estimate precisely, and we could only estimate an upper bound. Concerning the rate a we can argue as follows. First, recall that *a* = *α/δ*, where *α* is the rate at which spores make a transition from the GL into the AL. This rate can be written as *p_cross_/t_w_*, where is the average waiting time for a spore to enter a large eddy that takes it to the top of the ABL, and *p_cross_* is the probability of crossing the capping inversion once a spore is near it. Crossing of the capping inversion is a complicated process that is a subject of ongoing investigation, for example [57]. The likelihood to cross into the FA is highest in the evening hours when the ABL shrinks, leaving particles behind. However, even in the evening hours, most particles will remain in the ABL, so this probability is less than one. In other words, out of *N* spores that end up near the inversion layer, much less than *N* will cross it. Their most likely fate is to be taken back down to the ground by a large-scale eddy. But even if this probability was equal to 1, the upper bound on a is 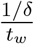. We estimated that 1/*δ* ~ 10 *s*. The average waiting time to enter a large-scale eddy is much larger than 10 seconds **cite**. Therefore, *a* ≪ 1.

Switching from AL to GL can happen via either a dry deposition process, or a a rain-down process. Rain-down events are episodic, and to incorporate such events would require *b* to be a time-dependent stochastic parameter. We briefly discuss the effect of time-dependent parameters in Appendix D, but it is not the focus of this paper. The likelihood of crossing into the ABL is highest in the morning hours, when the ABL swells. Following this penetration of spores into the ABL, deposition to the ground by gravity alone takes on the order of 1 day [58]. On the other hand, descending down on the same large-scale eddies that took spores up takes on the order of a half hour, and this is therefore the dominant descending mechanism. Therefore, *b* is expected to be of the same order as *a*. So, both are much less than 1 in our units.

The parameter 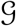 is hard to estimate. For this reason, all results in the paper were presented as a function of 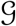. The reader can read off results that correspond to the value of 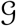 relevant to a specific pathogen of their interest – remembering that Δ*t*’ = 1 of dimensionless time corresponds to Δ*t* = 1/*δ* of physical time (which, for Wheat Stem Rust is ≈ 10 seconds, as argued above).

### A.1 Spore death rate due to UV radiation

UVB radiation is a potent killer of fungal spores, [26], [54], [55]. The curve representing the fraction of viable spores versus time has a shoulder, followed by an exponential decrease in time [26, 54]. Evidently, spores sense an accumulated dose of received radiation, such that a critical dose is required in order four UVB to start being affective. Aside from this memory effect, the exponential decrease in the number of viable spores indicates that spore death is a Poisson process during that stage. Therefore, we model spore death by a term such as —Δ*ρ* in our equations for density variables, where Δ is a spore death rate. In the absence of other processes, such as transport and new spore production, this leads to an exponential decrease in spore density – for example, *ρ* = *ρ*_0_*e*^-Δ*t*^.

Maddison and Manners [54] conducted a series of ground-based experiments with three species of rust spores; *Puccinia striiformis* was the most sensitive of the three. Based on their data, along with the shouldered functional form described above, and the model for altitude dependence, they made a series of predictions summarized as a graph of the fraction of surviving *P. striiformis* uredospores versus time for various times of the day, seasons, and altitudes. We extracted from this graph the decay rate at ground level, and at 3 km above ground in June. We found Δ to be 1.5 *hr*^-1^ at ground level and 3.2 *hr*^-1^ at 3 *km* above ground. These estimates refer to germinability. However, the relevant quantity is infectivity, which [54] estimate to be a factor of about 5 greater than sensitivity with respect to germinability. Thus, we must multiply the results by a factor of 5. Finally, converting to our dimensionless time, we get 2 × 10^-2^ at ground level and 4 × 10^-2^ at 3 *km* above ground, when rounding to 1 significant figure. The first of these numbers will represent *d_GL_*, and the second, *d_AL_* (although this is an over-estimate, since AL represents a combination of the effects of the lower parts of the free troposphere and the mean winds of the ABL). These are estimates for *P. striiformis* – one of the more UV sensitive of the Rust pathogens. Therefore this is an estimate for an upper range of spore death rates of Rust uredospores. For example, according to [54], *“P. graminis f.sp. tritici* race 21 uredospores are approximately 2-3 times more resistant to full sunlight than uredospores of *P. striiformis* race 2B”.

The values of Δ cited above for *P. striiformis* correspond to mean survival times of 0.7 hrs at ground level and 0.3 hrs at 3 km above ground. It would be useful to compare these estimates with spore death rates for other types of fugal crop pathogens. For this, we turn to Table 6.1 of Chapter 6 of Aylor’s book [26]. Aylor cites mean survival times for several types of pathogens. For example, sporangiospores of *Peronospora tabacina* – the pathogen that causes tobacco blue mold – will survive on average for 2.4 hrs under sunny conditions at ground level. Sporangiospores of *Pseudoperonospora cubensis* – the pathogen that causes downy mildew on cucurbits – will survive on average for 12 hrs under sunny conditions at ground level. The uredospores of *Uromyces appendiculatus* – another type of rust pathogen that attacks beans – will survive on average of 30 hrs (conditions not specified in [26]). The pathogen *Phytophthora infestans* – the potato late blight pathogen has a particularly small mean survival time of 1.2 hrs according to [26], but an even shorter time of 0.3 hrs is cited in [11].

All of these considerations suggest that the dimensionless rate for d on the order of 10^-2^ is actually an upper range for most species of pathogens. Some species will have *d* that are more than an order of magnitude below this. Moreover, the discussion so far concerned sunny conditions. The spore death rates under clouds can be orders of magnitude lower. For instance, Aylor [26] compares the numbers for mean survival times under sunny and cloudy conditions for two of the pathogens, and the lifetimes are a factor of 8 and 4. 4 larger. His 1986 paper [1] displays a dramatic decrease in the rate of spore death – many order of magnitudes – for a plume traveling under clouds versus under the open sun. Maddison and Manners in [54] provide data with similar conclusions. Therefore, under the cloudy conditions, the value of *d* can be several orders of magnitude below 10^-2^.

## B Saddle-point calculation of the front speed

We begin with linearized equations

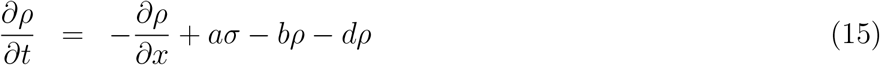

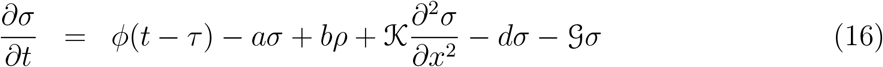

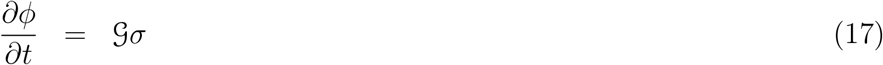

This has the form 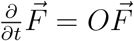, where

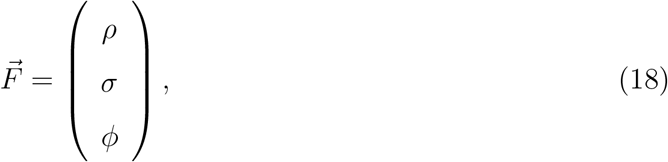

and the operator *O* has the form

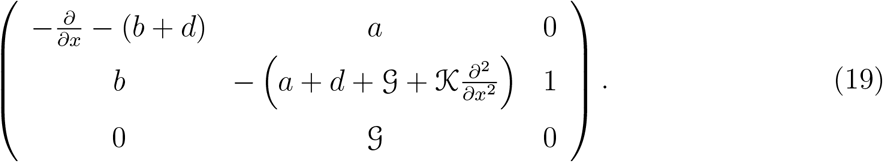

A general solution can be sought in the form of an eigenfunction expansion, 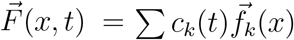 where 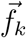 satisfies an eigenequation 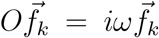 with boundary conditions. Here, the *i* is factored out in foresight of traveling-wave solutions, and the subscript *k* is just a label of the eigenfunction. Then 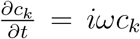, so the general solution has the form 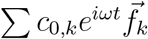.

The present problem is on an infinite domain, and it has free boundary conditions at ±∞. It follows that 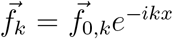. So, the label of the eigenfunction, *k*, is now seen to have the meaning of a wavevector. The general solution is just a superposition of traveling waves

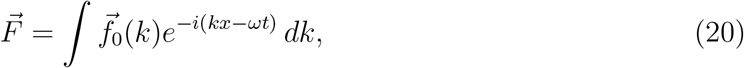

where 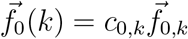, and we switched from the subscript to a functional notation because the set of 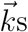 are continuous. Eq. (20) is a Fourier Transform solution to Eqs. (15)–(17). This is only possible because the problem has free boundaries. Then the Fourier basis ∝ *e^ikx^* are a valid (and complete) eigenbasis for this problem. If, on the other hand, the problem was on a finite domain with some boundary conditions on the ends, this basis generally would not satisfy such boundary conditions, and the set of 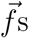 would be more complicated. In that case, the eigenfunction expansion would not take the form of a Fourier Transform.

Since the eigenvector 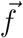 depends on the wavenumber *k* (*k* labels an eigenfunction), so does the eigenvalue, i.e. *ω* = *ω*(*k*).

### B.1 Saddle point method with zero delay, zero spore death, and zero diffusion

We first pursued the theory with zero delay (*τ* = 0), zero diffusion 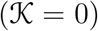, and zero spore death (*d* = 0). Even for this special case, the algebra is already quite unwieldy. The resulting dispersion relation satisfies the cubic equation

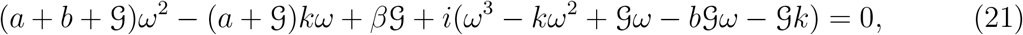

which can be obtained by direct substitution of 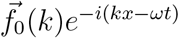 into Eqs. (15)–(17). This will also produce the eigenvector 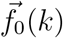, up to a multiplicative constant, which is set by the initial condition (IC). There are generally three branches, i.e. for each *k*, there is a triplet of eigenvalues and eigenvectors.

The idea of the saddle point method, as outlined in [33] is designed to find speeds of solutions such as Eq. (20), when they represent uniformly translating fronts, i.e. fronts that propagate with a constant speed, without changing shape. If the shape is not changing, then when we switch to a co-moving frame *z = x* — *s*_0_*t*, the solution will appear stationary. We evaluate the resulting integral via a saddle point *k*^*^. A more precise statement of stationarity is that the terms that are real and linear in t in the exponent of the integrand disappear (this does not eliminate all temporal variation of the solution, but it is designed to prevent an exponential growth of the solution). A reader is invited to [33] detailed explanation, which in turn is given in [42].

The mechanics of this procedure is as follows:

1. Find dispersion relation *ω*(*k*) by substituting 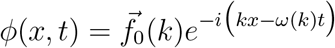 into Eqs. (15)–(17).
2. Find *k*\* from the solution to 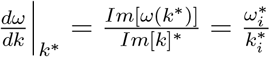.
3. Find speed 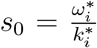. The spatial decay rate (for the downwind flank) or growth rate (for the upwind flank) rate can be shown to be 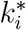.
4. Enforce 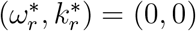 for non-oscillating solution.

Multiple solutions for *k*\* are possible, stemming from the fact that the method is not restricted to a specific IC, as long as the Fourier Transform of the IC is an entire function in *k*-space.

The saddle point method requires us to seek the wavevector *k*\* at which 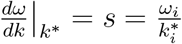. Applying this idea, and making use of implicit differentiation, would eventually lead us to the following expression:

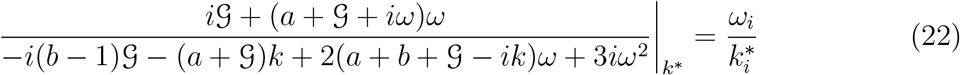

We set 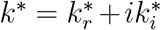, and 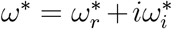, where *ω*\* = *ω*(*k*\*), i.e. dispersion relation evaluated at *k*^*^.

There are two ways to proceed. We can in principle solve for *ω*(*k*) from Eq. (21), substitute it into Eq. (22), solve for the corresponding complex saddle-point value of *k*\*, and finally, evaluate 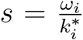. Alternatively, we can think of 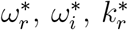, and 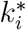 as four independent variables, and solve Eqs. (21)–(22) simultaneously (each has a real and imaginary part). Keeping in mind that 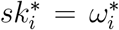, the variables to be solved for become 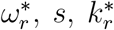, and 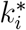. We will proceed this way. The actual calculation will be simplified by the fact that we have the additional criterion 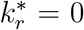 and 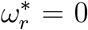, which physically avoids oscillatory solutions to prevent negative densities.

When this procedure is applied, we end up with the following pair of equations (we will drop the superscript “*” to lighten the notation):

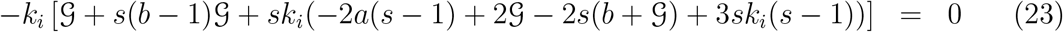

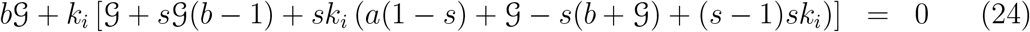

We then solve these for *s* as a function of parameters; we might also be interested in the decay rate of tails, *k_i_*, also as a function of parameters – the same two usual properties of fronts as discussed in [33] for the basic model. Note that there are multiple solutions to s as a function of parameters, and we will have to choose the correct solution based on limiting behaviors.

We performed a numerical solution of Eqs. (12)–(14) with *τ* set to 0, starting from a point initial condition (referred to as “simulations” *is that consistent?*). The speeds obtained from these simulations were then compared to predictions of the saddle-point theory. One of the roots matches the simulations, and this is shown in Fig. 18.

**Figure 18:**
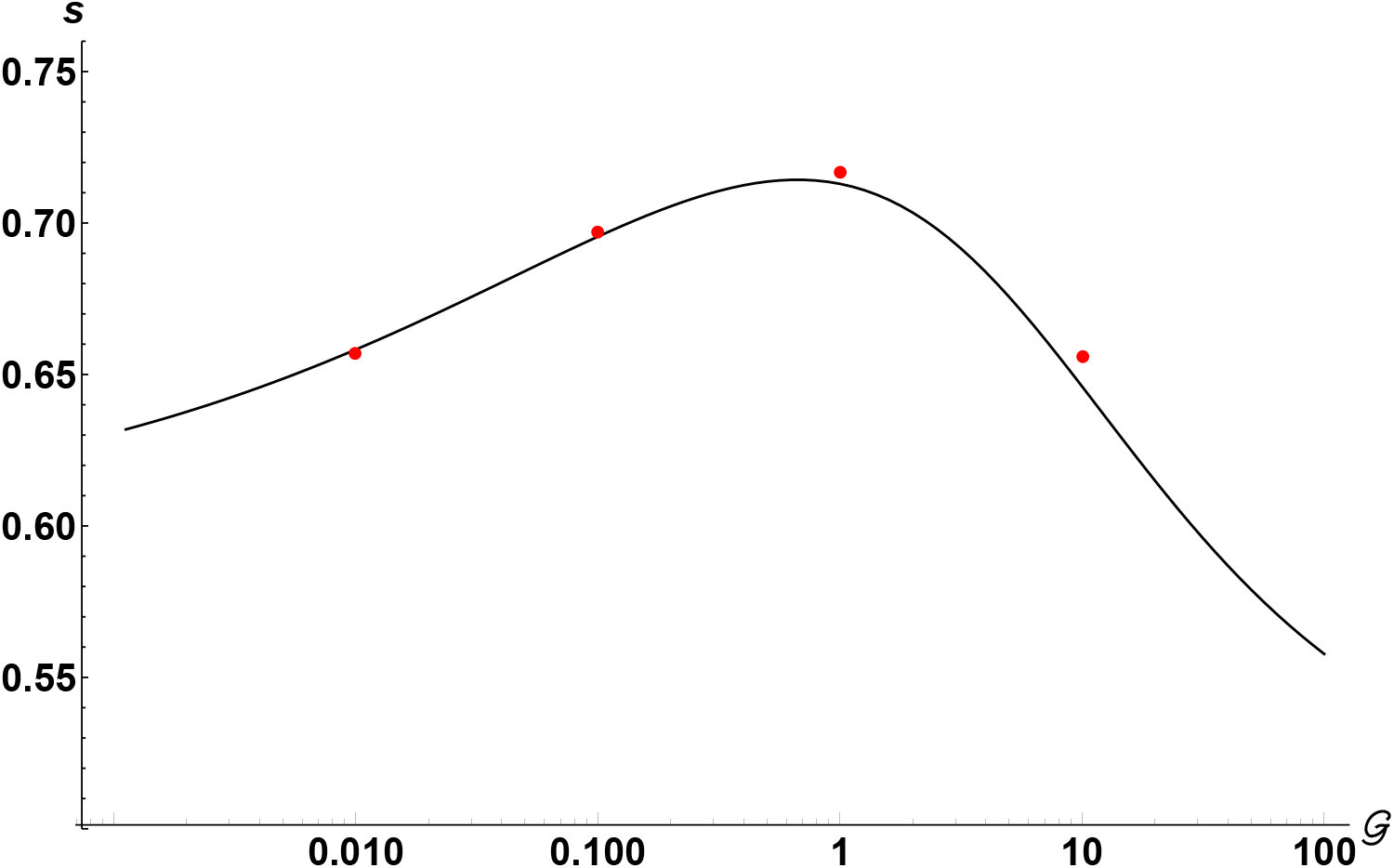
(Color online) Comparison of the speed of the leading front obtained from the numerical solution of Eqs. (12)–(14) with *τ* = 0 (red dots) with the saddle-point theory (solid line), generated by Eqs. (23)–(24).

To find the roots, we first solved for *k_i_* analytically from Eq. (23) – note that one of the roots is zero, and the other two roots obey a quadratic equation. The results were substituted into Eq. (24) and solved for s numerically. There are multiple roots. Some of them are complex. We can further constrain the real roots by noting that they have to approach 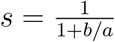 as 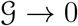, and 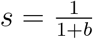 as 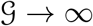. Out of all roots, we select only those that conform to these limits^1^. The first limit is easy to see from the observation that when 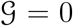, the model is equivalent to the original model with zero growth (and zero diffusion), which predicts a front speed of 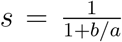 (that is, the advective speed times the fraction of the time spent advecting). The second limit can be obtained in the following way. Let’s divide Eqs. (23)–(24) by 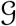

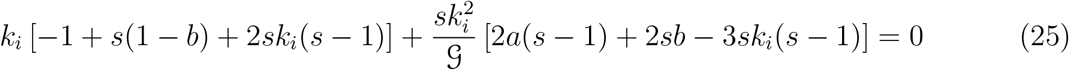

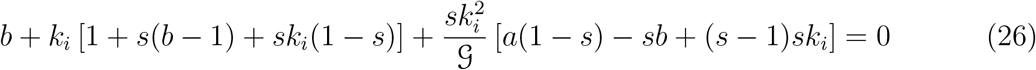

In the limit of 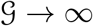 the 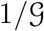 terms go away. The remaining terms can be solved for s and *k_i_*, giving
e

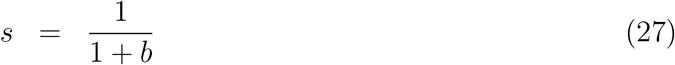

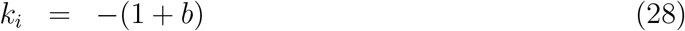

This procedure was applied for many values of 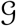. We start with an initial value of 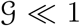, use an appropriate initial guess for *s* and *k*, and then feed the found value of *s* and *k* at the next, slightly larger value of 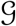. The comparison that is summarized in Fig. 18 suggests that the saddle-point theory works.

We can also extract the large 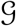 asymptotics (or tails) of 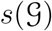. To do this, we seek a power law ansatz

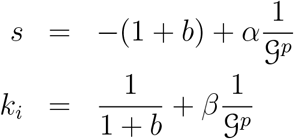

(we can think of this as a perturbation in 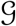 around zero). The power law that works (i.e. results match the full solution of 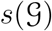) turns out to be *p* = 1/2. We substitute this ansatz into Eqs. (25)–(26), expand to lowest order in 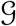, and solve for *α* and *β*. The result for the speed turns out to be

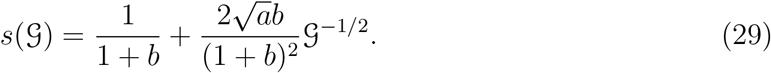

### B.2 Saddle point method with non-zero delay, zero spore death, and zero diffusion

We now add the delay. Thus, we will be studying the front speeds of the following linearized model:

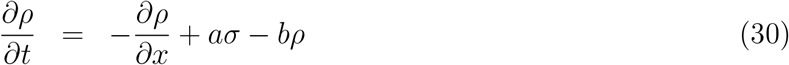

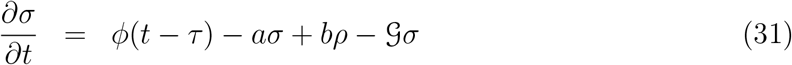

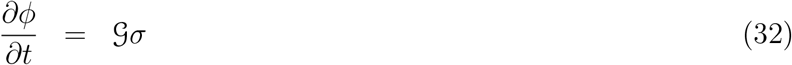

The new dispersion relation now obeys

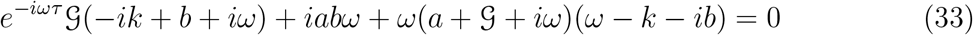

Note the presence of the term *e^-iωτ^*, which makes these equations transcendental. Following the same steps as before, we obtain – after setting 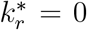 and 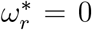 – the following two equations

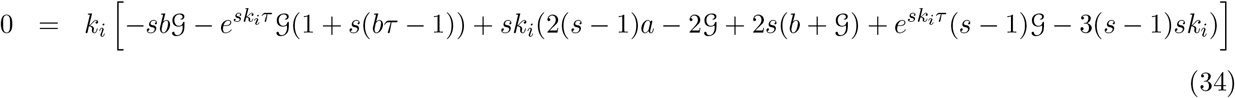

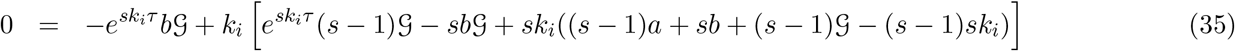

These transcendental equations are the equivalent of Eqs. (23)–(24). They are solved numerically, and we were able to produce saddle-point predictions of *s* vs. arbitrary 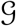 and *τ*. We compare the analytical predictions produced by the saddle-point method with predictions of speeds extracted from direct simulations of Eqs. (12)–(14). This is shown in Fig. 19 for a particular value of *τ* = 10.

**Figure 19:**
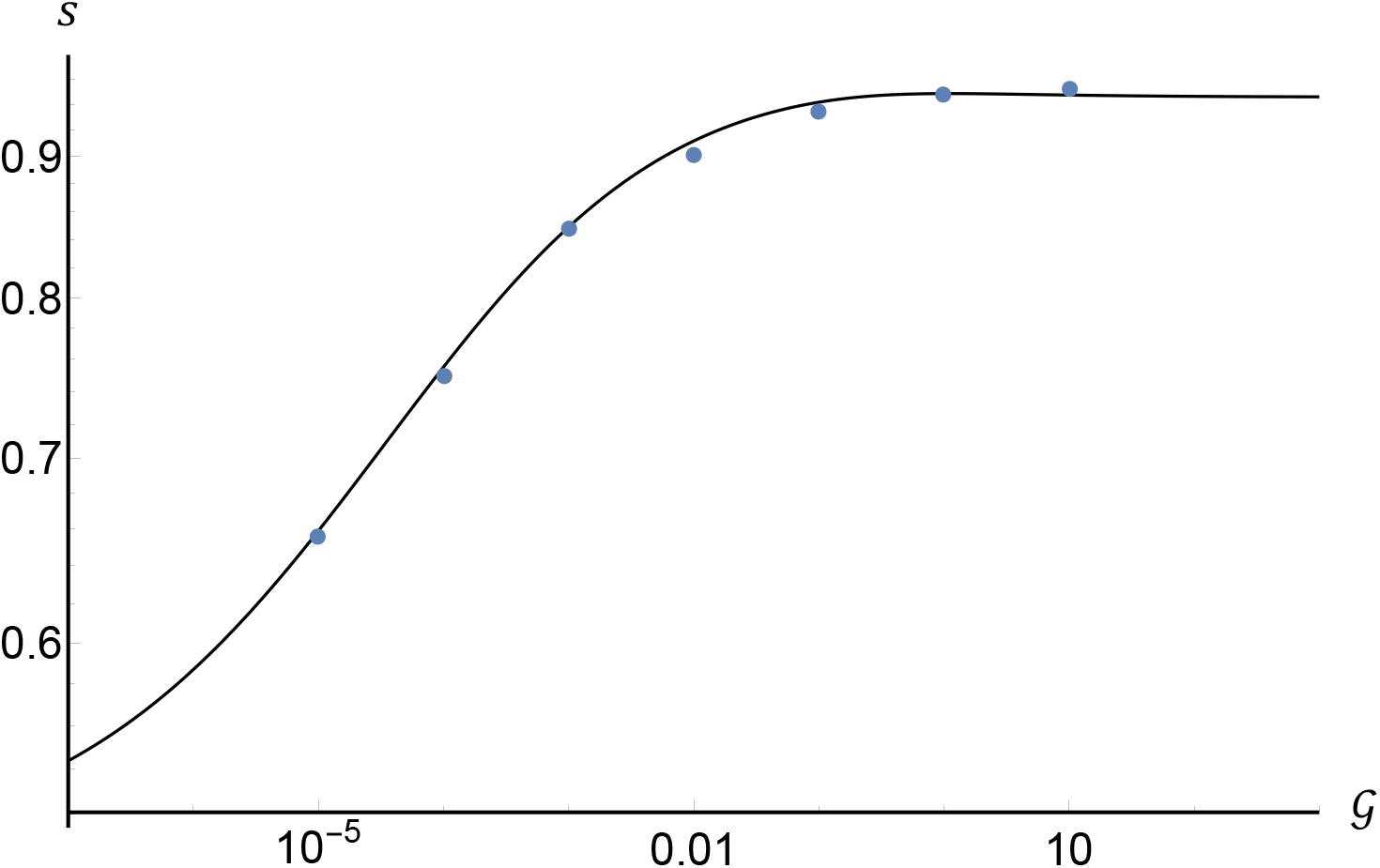
(Color online) Comparison of the speed of the leading front obtained from the numerical solution of Eqs. (12)–(14) (dots), with the saddle-point theory generated by Eqs. (34)–(35) (solid curve). Here *a* = *b* = 0.01, *τ* =10, *d* = 0, 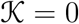.

The realistic values of *τ* in our units are of the order of 10^5^ (see Appendix A). It is in a situation like this, where theory is indispensable, since such large values of *τ* are inaccessible in simulations, due to dramatic increase in simulation time with *τ*. Fig. 20 demonstrates the change of 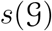 with increasing *τ*.

**Figure 20:**
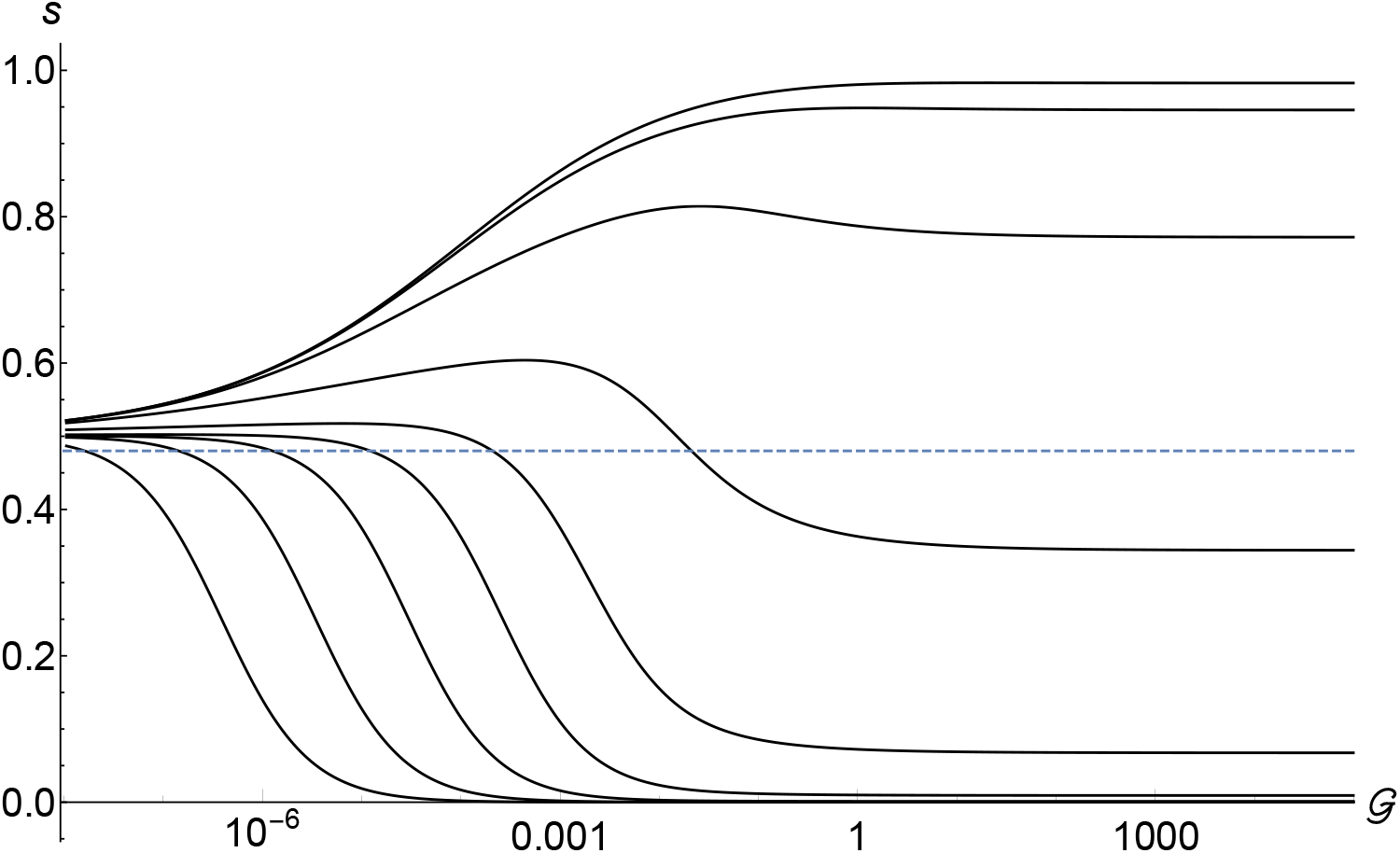
Predictions of the saddle-point theory at non-zero delay. The curves correspond to *τ* =1 (top), through 10^8^ (bottom) with increments by a factor of 10. Here *a* = *b* = 0.01.

We can again extract the form of tails of 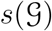 following a similar, albeit more involved, procedure as in Section B.1. The result is

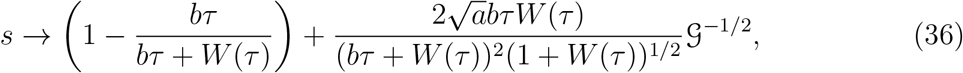

where *W*(*τ*) is a *w* that satisfies *τ* = *we^w^*, i.e. it is the inverse of the function *f* (*w*) = *we^w^*, also known as a Product Log or Lambert W function. Thus, 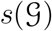 tends to 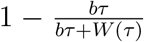 as 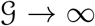. Subtracting this limiting value, we can display 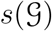 on a log-log plot, see Fig. 21. This is useful because it shows that there are actually *two* crossovers; the intermediate regime is clearly visible in Fig. 21. Therefore, extracting the value of the first (lower) crossover analytically by equating the large-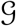 tail with the low-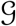 tail would give a wrong prediction for the scaling of the lower crossover point. However, we can easily extract the scaling of that lower crossover point by directly measuring the value of the function at a characteristic contour value where the crossover begins (dashed line in Fig. 20). The data collected this way is depicted in Fig. 22. It is fit very well by a function 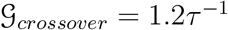.

**Figure 21:**
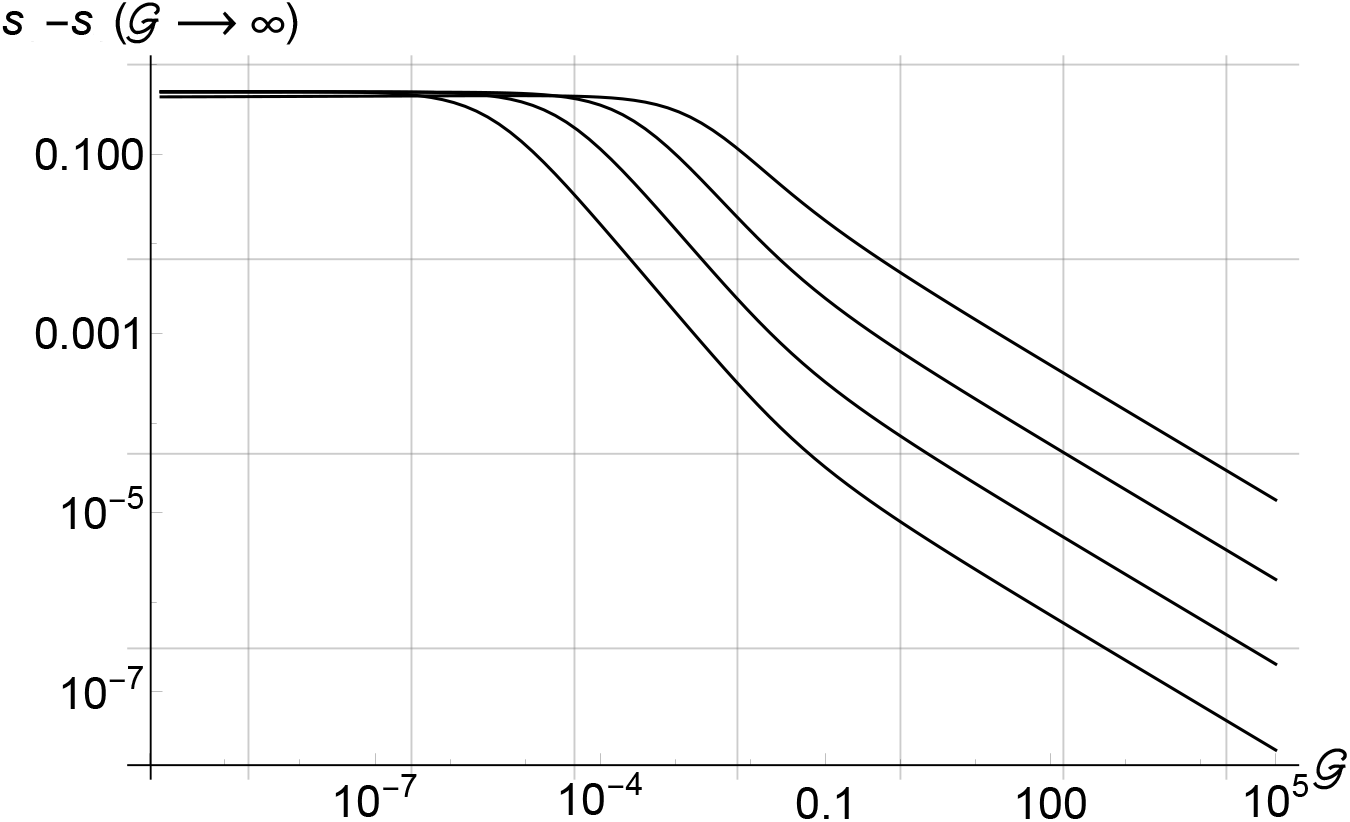
Same as Fig. 20 on a log-log plot. This figure shows the existence of two crossovers. Here the *τ* values range from 10^4^ to 10^7^ in steps of 10.

**Figure 22:**
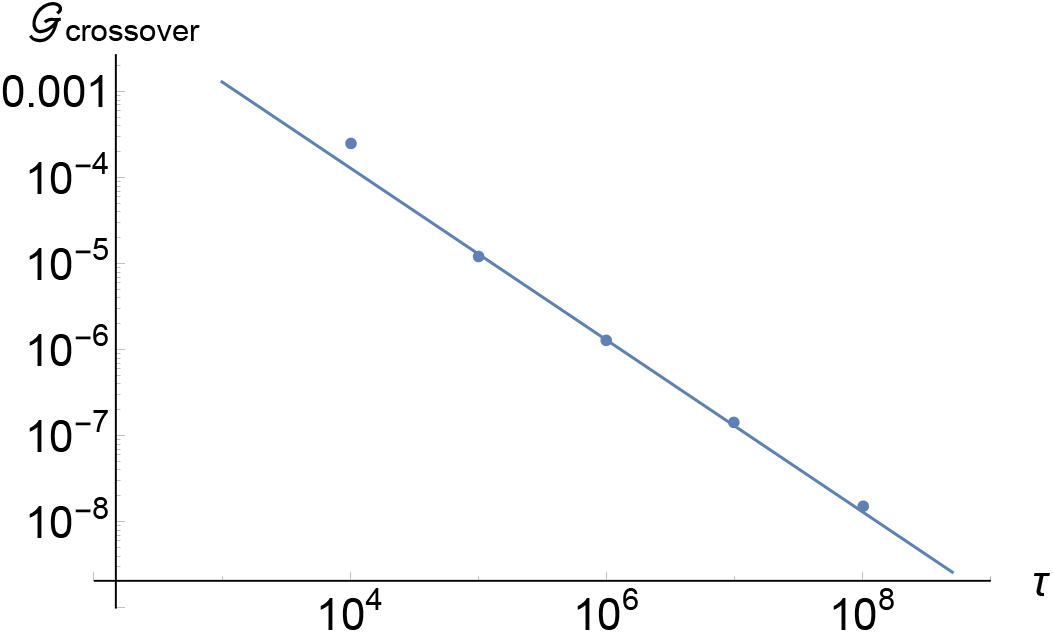
(Color online) 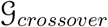 vs. *τ*. We can see the beginning of the departure from the power law at lower *τ*.

This scaling only applies to the high-*τ* tail; for lower *τ* the data lies above this trend. This is clearly seen in Fig. 20 (note the intersection of the graphs with the dotted reference line). In Appendix A, we estimate that *τ* ranges in the hundreds of thousands in our units, suggesting that 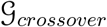 would be of the order of 10^-5^. As long as 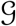 is above this very small value, the latency will slow down the front.

The simple scaling 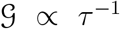 suggests a simple physical interpretation of the lower crossover. There are two three time scales (and associated rates) that control the spore production rate. The first is the rate at which uredinia produce spores. This rate is *δ* in physical units, but it becomes 1 after non-dimensionalization, so it is set to 1 without loss of generality. Thus, the associated time is also 1. The second is the rate at which GL spores infect host plants. That rate is 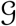, and thus 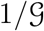 is an average waiting time for a spore to infect a plant. The third time scale is *τ* – the waiting time between infection and production of new spores. When 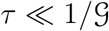, the dominant waiting time is the waiting time to infect the plant, and latency is not expected to play a role; on the time scale of 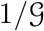, the time spent within the plant is negligible. When 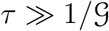, the dominant waiting time is the latent time – latency should generally affect the front speed in this regime. Thus, the crossover between regime in which the speed will not depend on *τ*, and when it will depend on *τ* is expected to be around 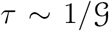 or 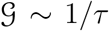. This is indeed what is happening. When both *τ* and 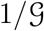 are much larger than the third timescale – 1 in our units – these two will determine the crossover.

We estimate in Appendix A that for diseases such as wheat stem rust with a latent period of about two weeks, *τ* is of the order of 10^5^. The parameters *a, b* are both expected to be ≪ 1. We plot multiple *s* vs. 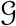 curves for this parameter range in Fig. 23. We see that the crossover as defined above, takes place at 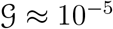.

**Figure 23:**
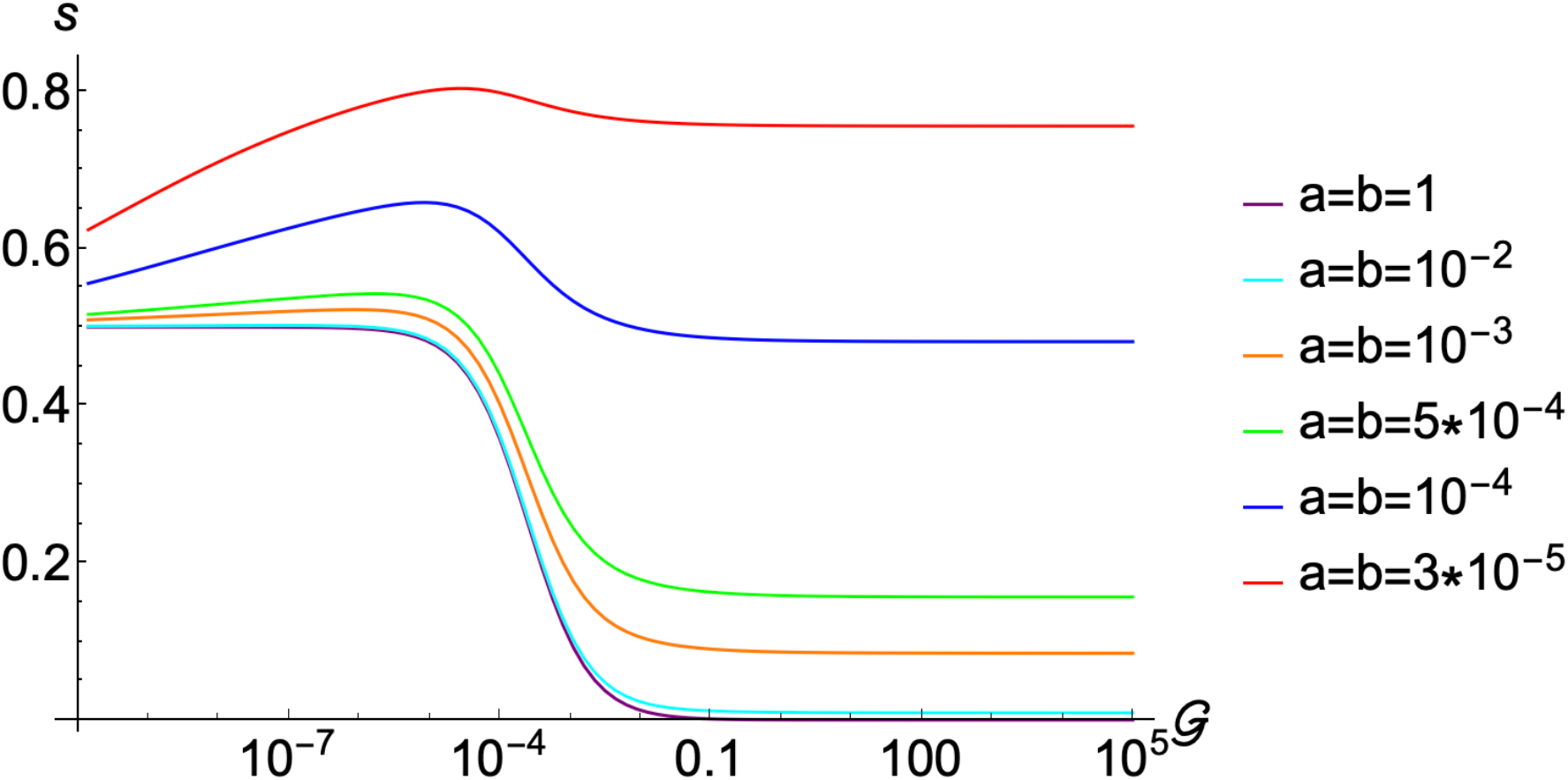
(Color online) *τ* = 10^5^. The six curves, from lowest to highest are *a = b* =1 (bottom), 10^-2^, 10^-3^, 5 × 10^-4^, 10^-4^, 3 × 10^-5^ (top).

The tail in the large-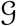 regime has a value given by the formula 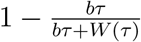. It goes to 1 as *b* → 0, and goes to zero at large *b*. The tail in the small-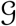 regime is given by 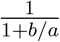 – the speed of the front in the basic model with zero growth.

### B.3 Saddle point method for the full model

We finally apply the saddle-point method to the whole set of Eqs. (15)–(17), with all the parameters being non-zero. The resulting equations are:

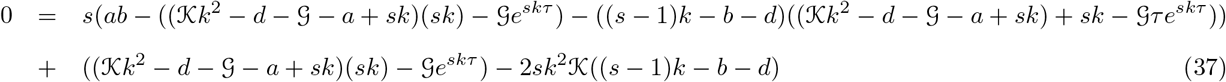

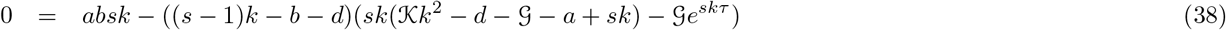

We again solve them numerically. The results presented in Section 3.1 were based on these numerical solutions.

It is important to emphasize that equations such as these give a set of possible solutions, but they do not address the question of solution selection. In general, Eqs. (37)–(38) can predict many solutions – three or even more pairs of (*s, k*). In other words, each of these equations identifies a curve in (*s, k*), space, which may intersect at several points. Which solution (or which intersection) will actually be selected by the initial condition is an additional problem – one for which we do not have a theory. However, we have two additional constraints that taught us which intersection corresponds to correct predictions for the selected *s*. The first is a check against simulations. The second is a limit of 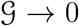 at zero *d*. We must choose such an intersection that in this limit gives 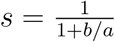 – the result of a basic model with zero growth. These two checks taught us which intersection to pick.

We will now illustrate our method. Consider parameters depicted in Fig. 5: *τ* = 1, 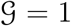, *d* = 0. The following is a series of portraits of (*s, k*) plane at progressively larger 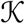. In Fig. 24 we show (*s,k*) for *a* = *b* = 0.01. At least three solutions (intersections) are seen. The solution that matches the simulations is indicated by an open red circle. To see this, we compare this solution with simulations in Fig. 26. On the other hand, when *a = b* = 0, the contours in (*s, k*) plane are different – these are shown in Fig. 25. The solution that matches the simulations is indicated by a filled blue circle. To see this, we compare this solution with simulations in Fig. 27. In situations such as the case with *a = b* = 0, when roots cross, we have to compute the theoretical curve by pieces, in order to track the correct root. However, we have not encountered this scenario when studying non-zero *a = b* cases.

**Figure 24:**
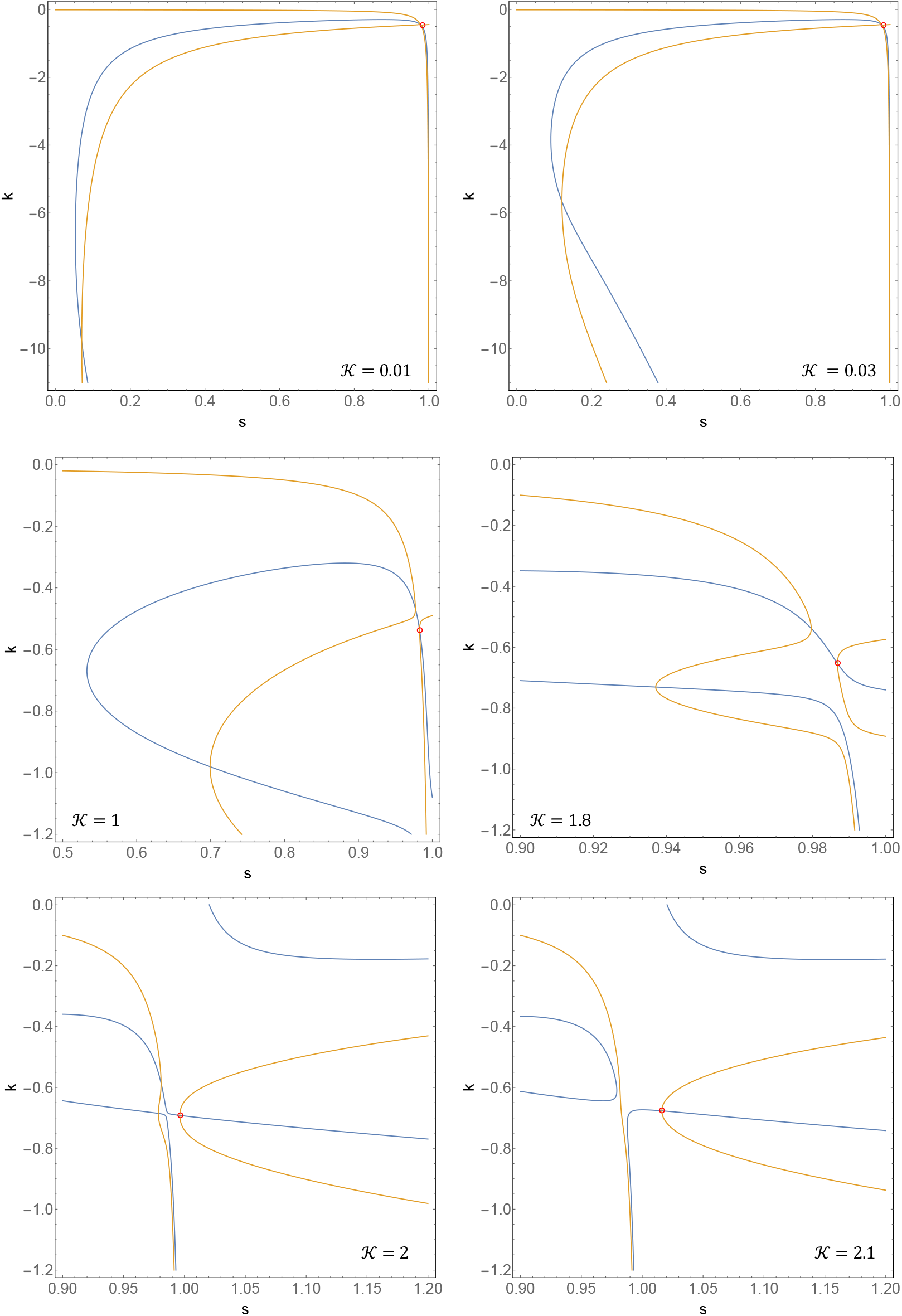
(Color online) (*s, k*) plane at different values ok 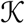. Here *a = b* = 0.01. The blue curve represents the contour corresponding to Eq. (37), while the orange curve represents the contour corresponding to Eq. (38).

**Figure 25:**
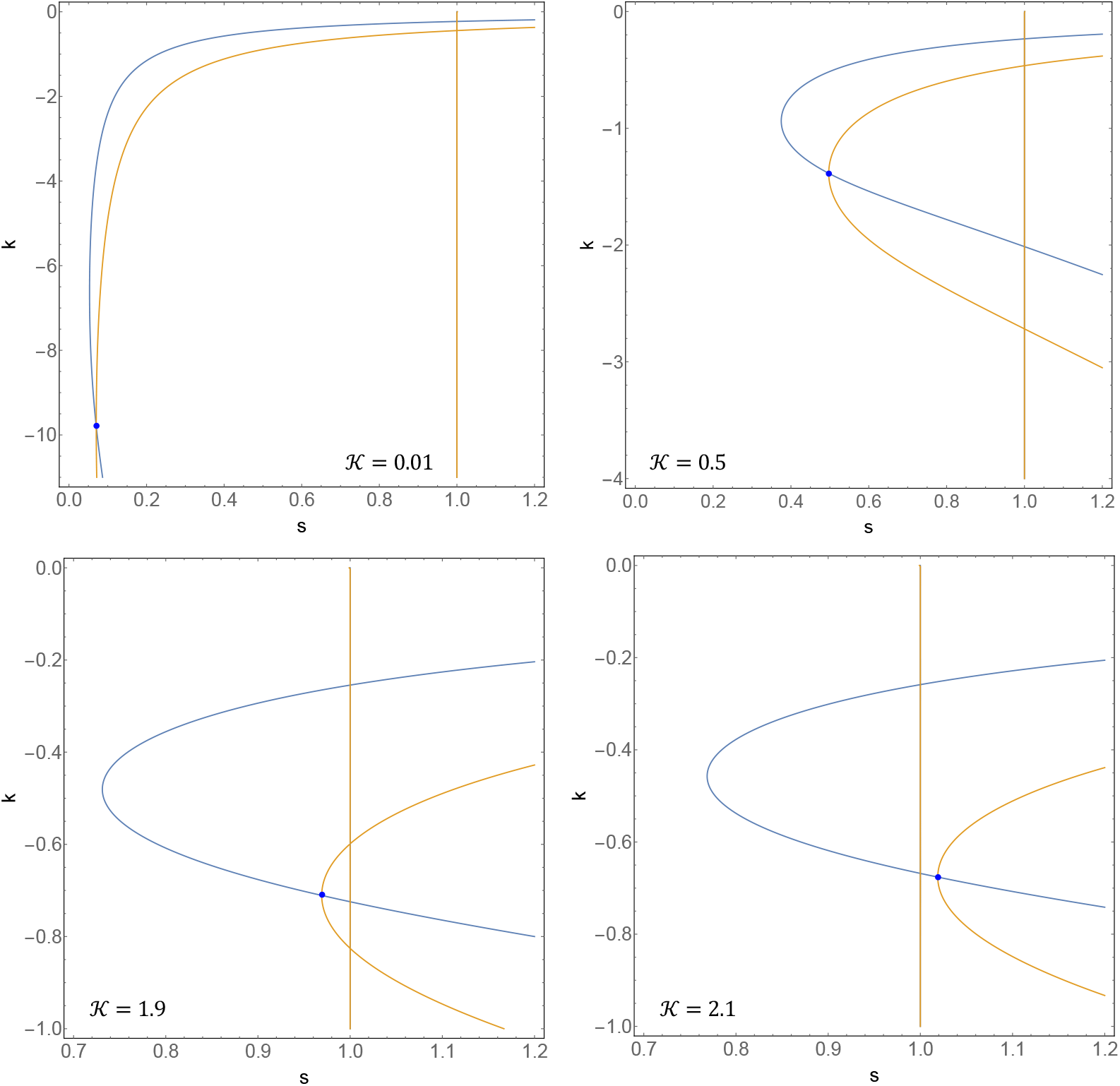
(Color online) (*s,k*) plane at different values ok 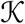. Here *a = b* = 0. The blue curve represents the contour corresponding to Eq. (37), while the orange curve represents the contour corresponding to Eq. (38).

**Figure 26:**
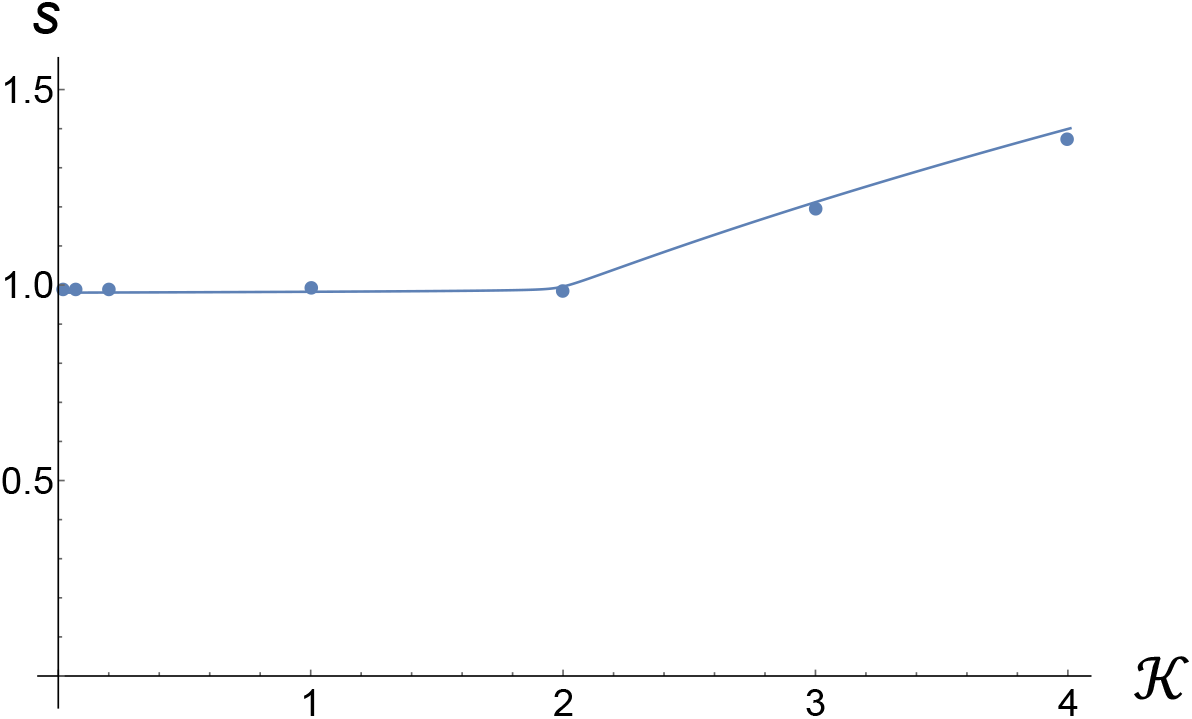
(Color online) Comparison of simulation data (dots) with analytical predictions based on the root from Fig. 24 denoted by a red circle. This is part of the same data that is presented in 5.

**Figure 27:**
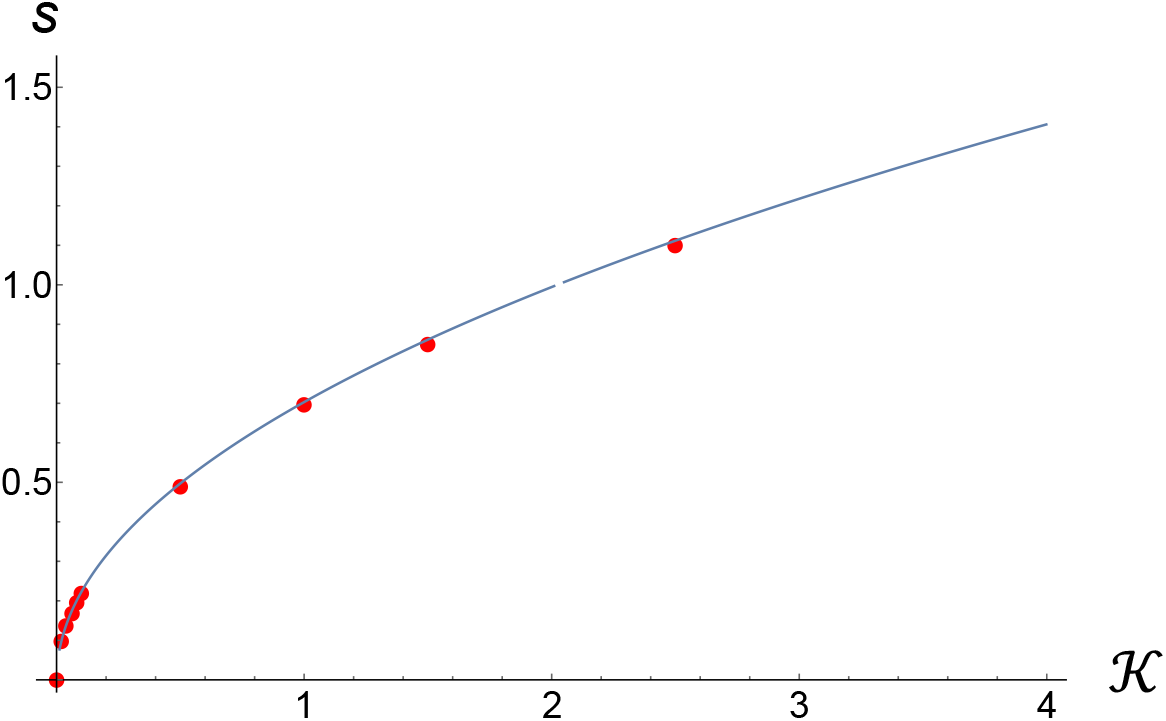
(Color online) Comparison of simulation data (dots) with analytical predictions based on the root from Fig. 25 denoted by a solid blue dot. This is part of the same data that is presented in 5.

In situations when simulations are not available – most notably at large *τ*, we also used another constraint – that we are interested in such a root that evolves to 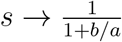 as 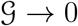 at zero *d*. We found that the branch structure in (*s,k*) space evolves continuously as d is made non-zero. In addition, in the situations analyzed in this paper – particularly the ones described in Section 3.1, the branch structure in (*s, k*) space always had the same topology as the one depicted in Fig. 24, and the circled root was the one that gave 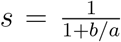 as 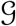 approached zero and *d* was set to zero.

### B.4 Different spore death rates

When the spore death rates in the AL and GL are different, and are respectively give by *d_AL_* and *d_GL_* Eqs. (37) and (38) generalize to

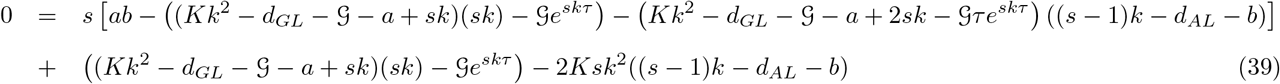

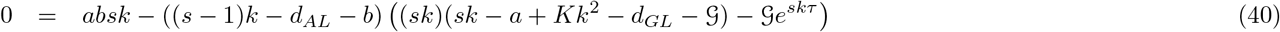

When *d_AL_* → ∞, the two equations reduce to

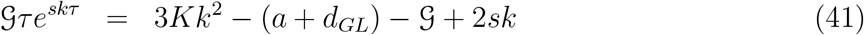

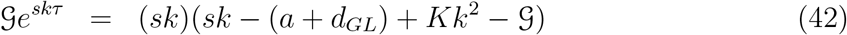

We note that parameter b dropped out from these equations. Also note that *a* and *d_GL_* appear additively.

## C Some additional useful results

The following study probes the importance of spore death rate. In Fig. 28, we plot speed of the coupled branch at 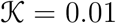 versus the spore death rate *d*, while holding the value of 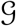 fixed; c.f. Fig. 9. We see that past some critical *d*, the gap decreases as a power law.

**Figure 28:**
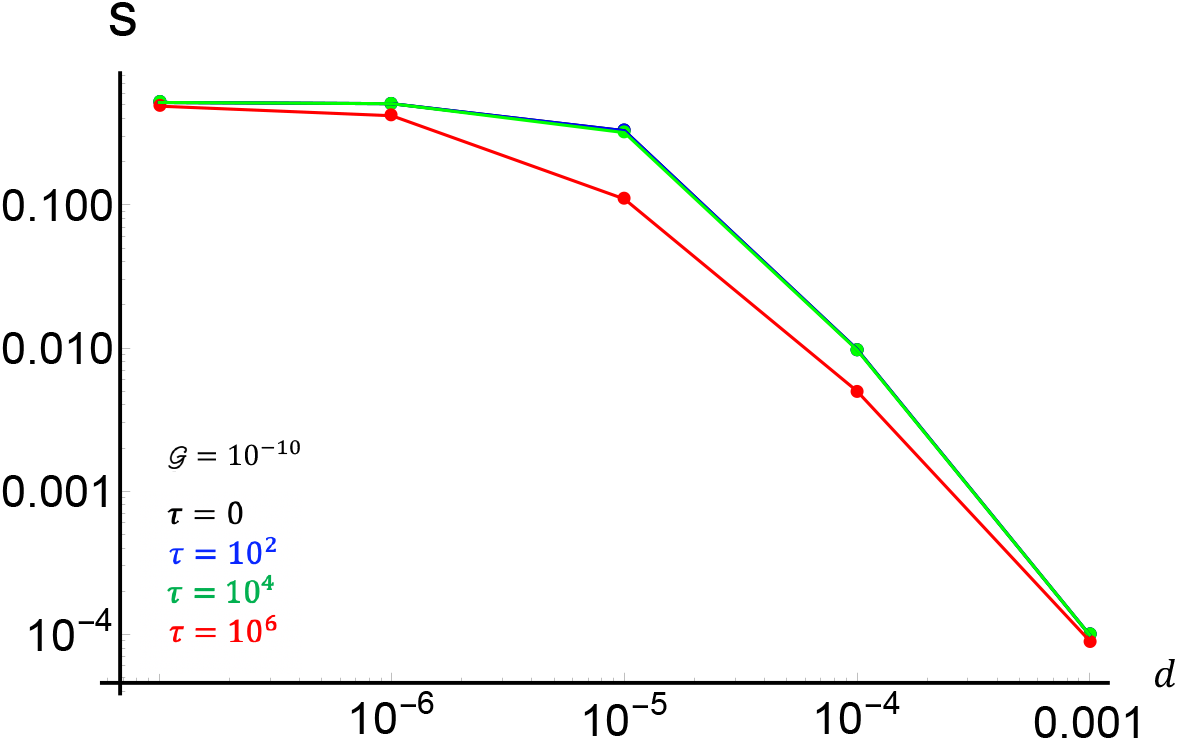
(Color online)Gap size versus *d* at a fixed 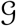 for several values of latent times, *τ*.

The importance of spore death can be elucidated even further by directly plotting the speed of the front versus the spore death rate. Fig. 29 displays the front speed vs. the spore death rate *d* for two values of 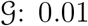 and 1. The tail of the 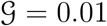 data scales like *d*^-2^.

**Figure 29:**
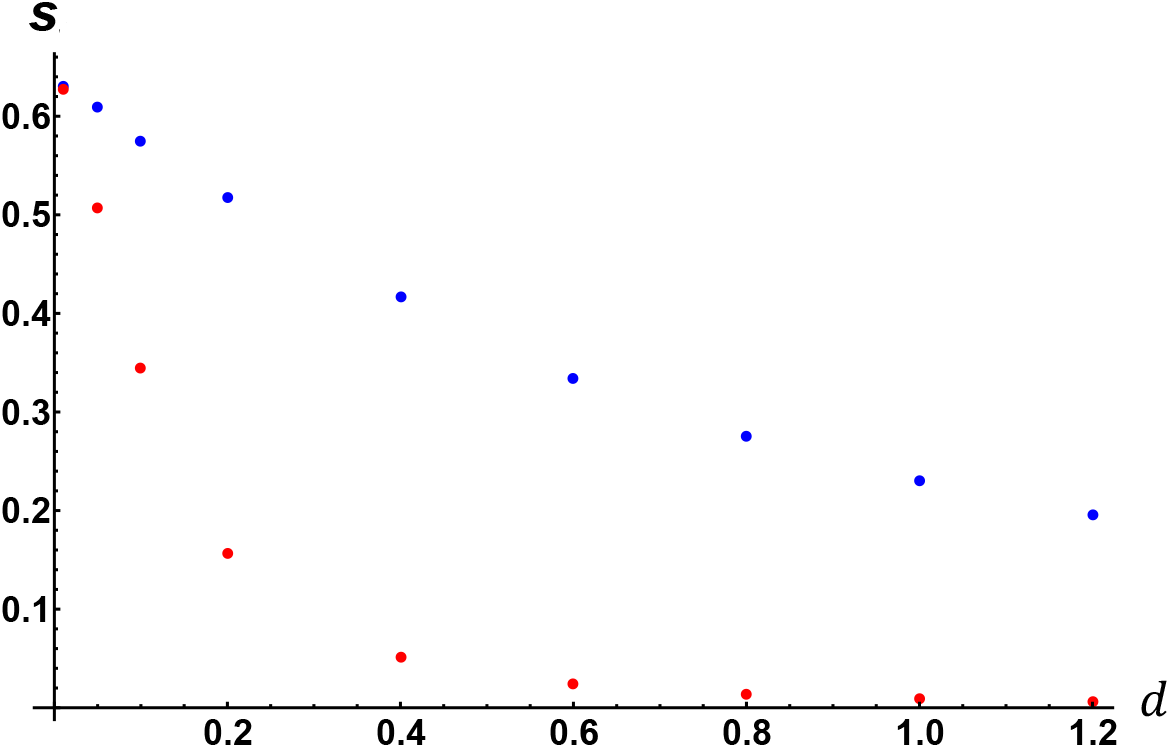
(Color online) *s* vs. *d* at *a* =1.5, *b* =1. The two values of 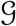 are 1 (blue, upper curve) and 0.01 (red, lower curve). Here 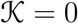.

In the absence of the spore death, and with only the latency effect, the speed approaches ≈ 0.63, whereas in the absence of both the spore death and the delay, the speed would be ≈ 0.83 (see the formula in the caption to Fig. 3). The faster front at larger 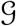 is due to the fact that spore death does not affect the fungal state. Thus, if spores can transition to this state sooner, this biomass is preserved at a larger rate. On the other hand, having a small S dramatically decreases the front speed, as it forces the spores to be exposed to UV light longer, providing only a rare opportunity to transition into the fungal state.

## D Other study: time dependence of parameters

We considered the role of temporal variation of parameters in the basic model, Eqs. (5)–(6).

### D.1 Varying coupling rates

For this numerical experiment, we set 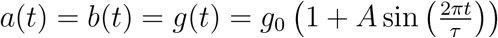. The diffusion coefficient 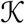 was set to zero, and we used *g*_0_ = 0.1. We explored the (*A, τ*) space. First, it deserves to note that when variation is sufficiently slow (*τ* is sufficiently large), the parameters become quasi-static, and the front speed is set by the instantaneous value of *g*

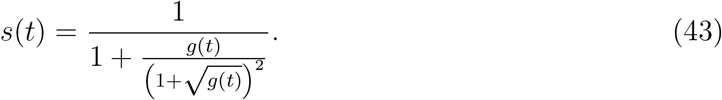

In other words, on the time scale of the variation, the front is able to adjust its speed instantaneously. We will call this “adiabatic theory”. This will be true whenever the rate of the temporal variation of parameters is much smaller than the relaxation rate of the model with constant parameters; that’s the meaning of “sufficiently slow”. We don’t know exactly this relaxation rate (it may be comparable to the transient time in the time-independent model, for which we do not have a theory), but we will see that as *τ* is made progressively greater, the adiabatic prediction of Eq. (43) will become progressively more accurate.

The opposite limit is very small *τ*, where again, the smallness is measured in comparison to the relaxation time. In this regime, the results will probably not be dependent on the amplitude of the variation, since the system will only be able to “feel” the average *g*.

Although we do not have a theory for the value of the relaxation time, dimensional reasoning suggests that it can not exceed 1 by more than an order of magnitude, since all the parameters in the model are of order 1 – so presumably, a temporal parameter that depends on these order-unity parameters will also be of a similar order – although this is not guaranteed.

Since the variation of *g* was periodic, the response of the speed was also periodic – it had a maximal and the minimal value. We will plot this speed below, but first, we plot the max, min, and the average value of the speed as a function of the amplitude of variation when *τ* = 100. Based on the reasoning in the previous paragraph we expected this to be in the adiabatic regime. These are plotted below for both the *σ* and the *ρ* densities.

The adiabatic predictions are given by

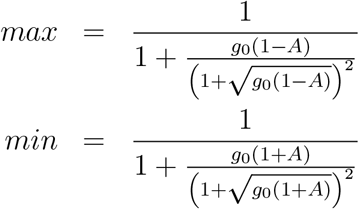

Several features of these graphs deserve discussion. First, we note that the adiabatic simulation data does follow the adiabatic prediction very closely (note the scale on the vertical axes). The slight spread in the points as *A* → 0 is a function of the finite step size in the numerical solution of PDEs; we verified that it decreases with a degreasing step size. A striking departure from the adiabatic trend is obvious in the right panel at *A* = 1.

We plot a portion of the results of speed vs. time corresponding to these parameters in Fig. 31. This data also shows very sharp spikes in the speed – which are the cause of anomalously large maxima and small minima. However, what’s especially peculiar is that the speed on the Ground Layer exceeds the wind speed (1) during these short episodes. How is this possible? Note that the advective speed is bound by the wind speed. Also note that here *A* = 1, which means that for an instant during a cycle, the exchange of matter between the two layers is shut off. During a short window of time when this happens, the advective currents continue to move the material in the downwind direction. When the coupling begins to increase back from zero, the spores rain down on a clean segment of the ground *all at once,* leading to a very large speed of the growth front. The speed is not infinite because there is a gradient in the AL density, leading to a gradient in the density of spores that fall on this clean patch of GL. Thus, the anomalously large speed is the speed of the *growth front.* Such anomalous spikes require A to be close to 1 to accommodate this “shut down” of the coupling between the two layers during a short portion of the cycle.

**Figure 30:**
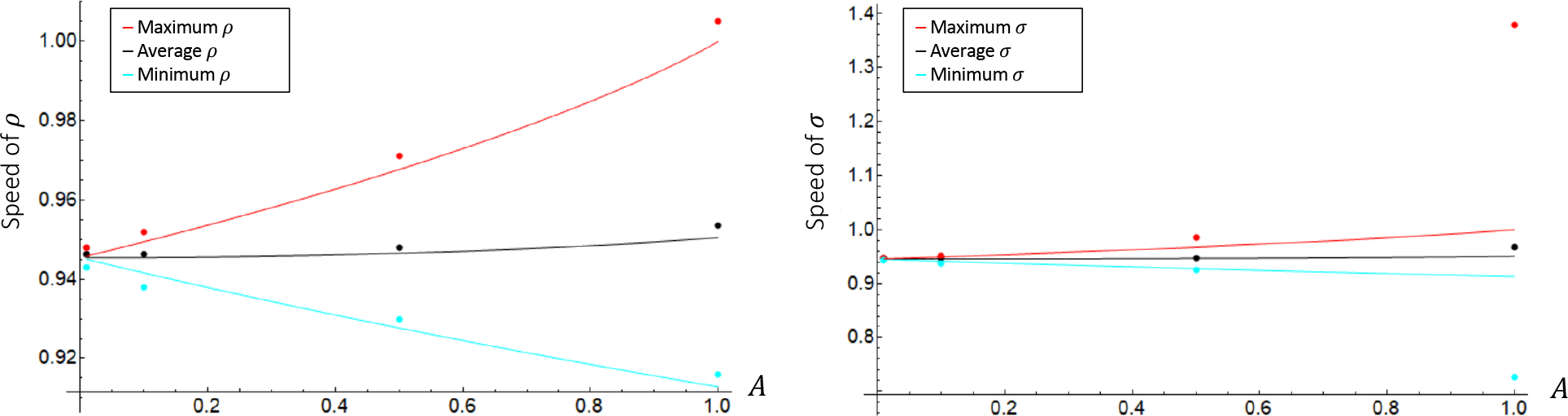
(Color online) Min, max, and average of the variation in the speed of each front. Dots – simulations, solid curves – adiabatic theory. The speeds are in units of advective speed. Here *τ* = 100. All the curves approach 0.945, which is the speed of the unmodulated front at these *a* and *b*. Note also that this is not exactly the same as the average of the modulated speed – whether from simulations or even from the adiabatic theory.

**Figure 31:**
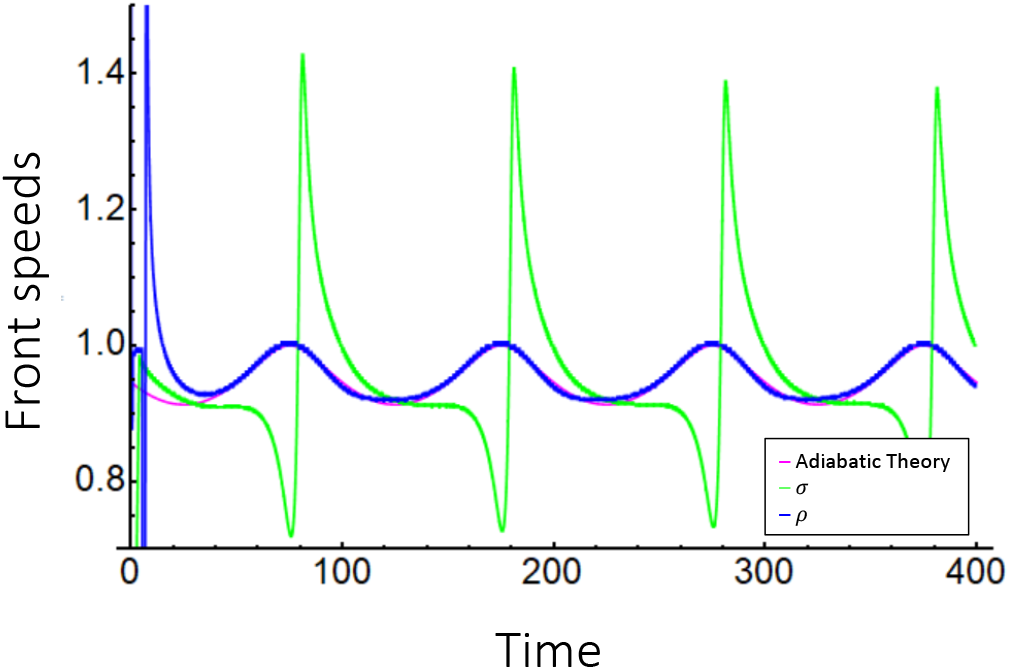
(Color online) Front speeds (in units of advective speed) versus time (in units of 1/*δ* of the basic model, which in conventional units is about 3 hours – see Sec. ??). Here *A* =1.

As the period of the variation of *g* decreases, the results begin to fall off the predictions of the adiabatic theory. This is show in the next two graphs.

**Figure 32:**
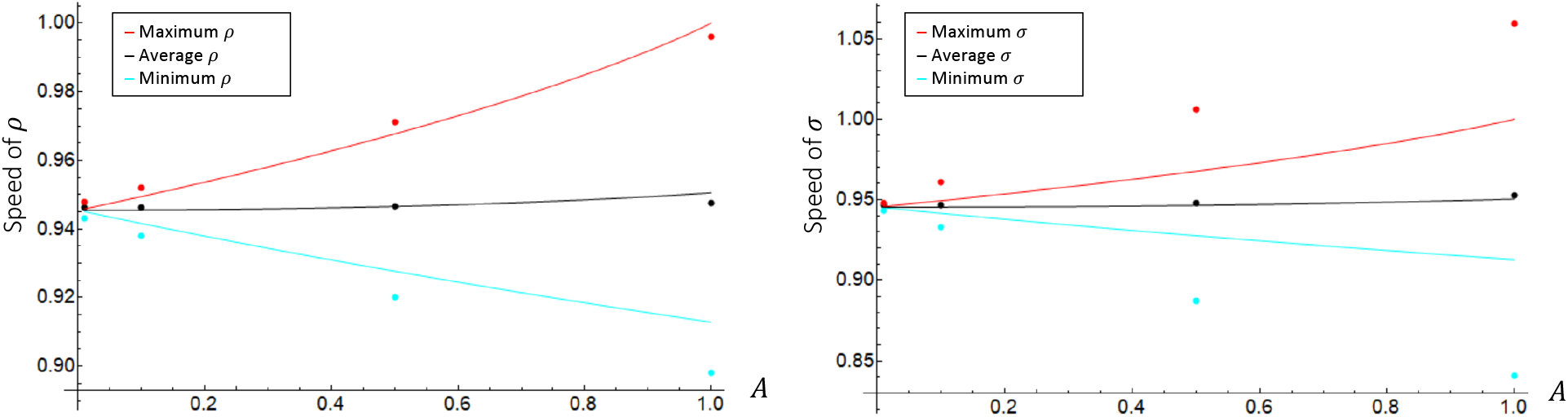
(Color online) Min, max, and average of the variation in the speed of each front. Dots – simulations, solid curves – adiabatic theory. The speeds are in units of advective speed. Here *τ* = 1. All the curves approach 0.945, which is the speed of the unmodulated front at these *a* and *b*

**Figure 33:**
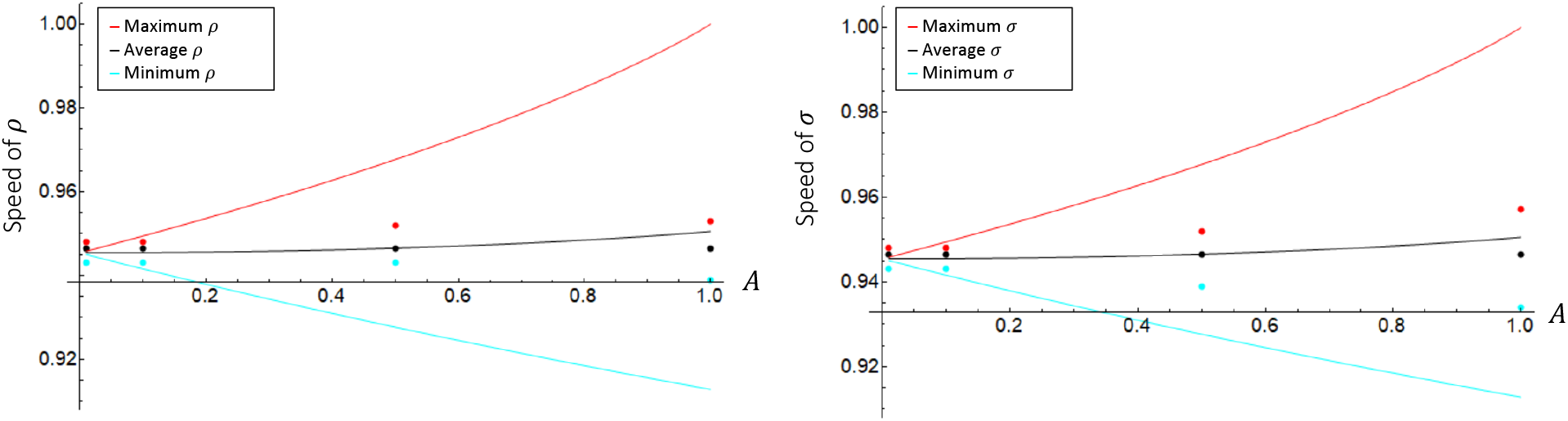
(Color online) Min, max, and average of the variation in the speed of each front. Dots – simulations, solid curves – adiabatic theory. The speeds are in units of advective speed. Here *τ* = 0.1. All the curves approach 0.945, which is the speed of the unmodulated front at these *a* and *b*

The corresponding speed vs. time graphs are shown next.

**Figure 34:**
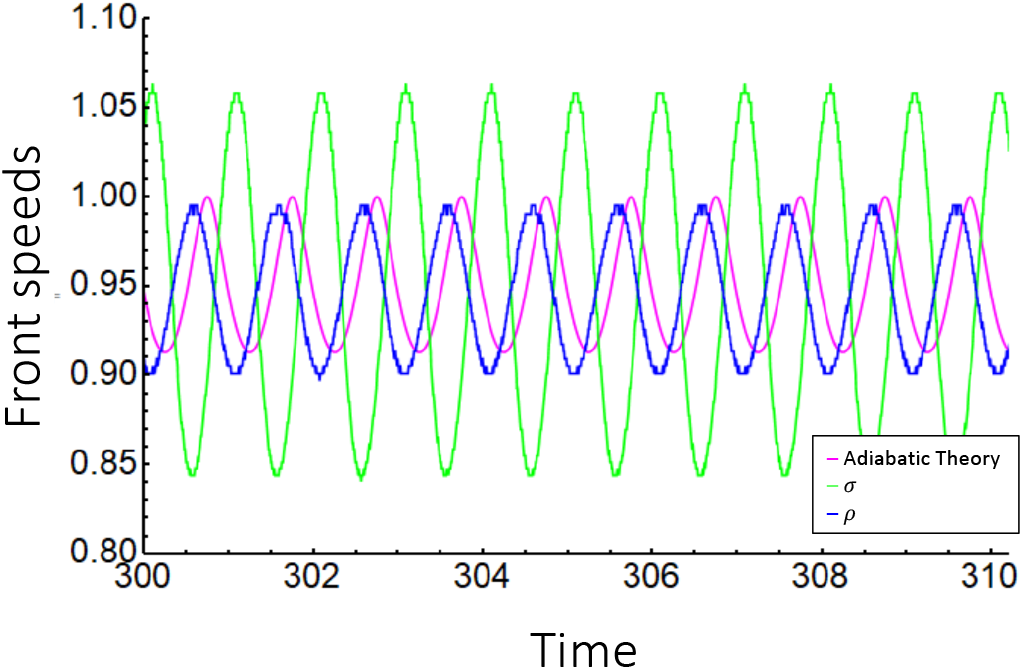
(Color online) Front speeds (in units of advective speed) versus time (in units of 1/*δ* of the basic model). Here *A* = 1.

**Figure 35:**
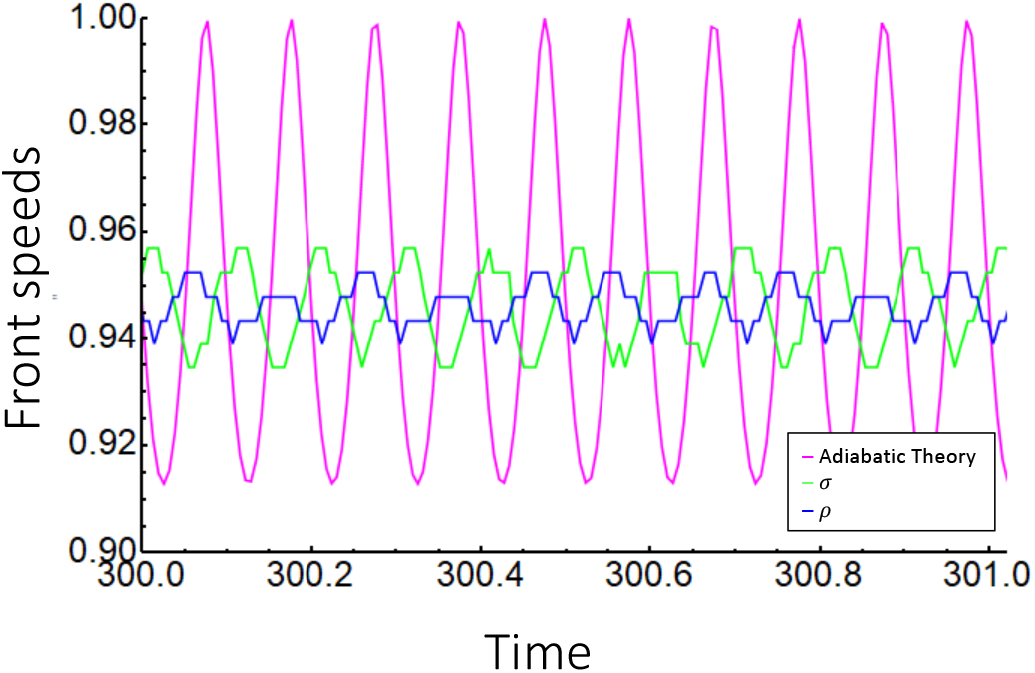
(Color online) Front speeds (in units of advective speed) versus time (in units of 1/*δ* of the basic model). Here *A* = 1.

Evidently, there is a phenomenon similar to parametric resonance [56], where the amplitude and phase of the speed variation is a function of the frequency of the variation of parameters. We summarize the amplitude (here defined as *max — min*) dependence upon the period of the parameter variation in Fig. 36.

**Figure 36:**
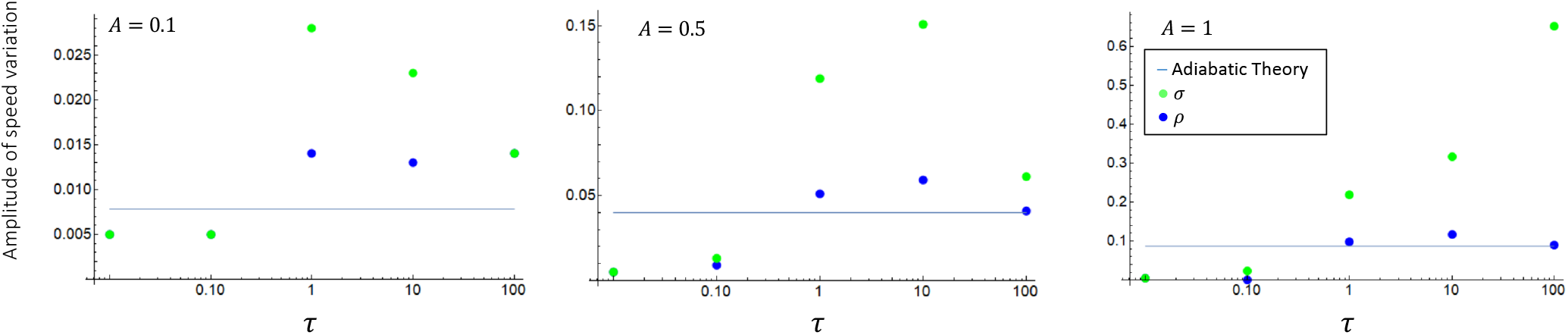
(Color online) Amplitude of speed variation vs. the period *τ* of the variation of *g*(*t*). Some green and blue dots coincide.

The time *τ* is in units of 1/*δ* of the basic model. As discussed in Sec. ??, 1/*δ* ≈ 3 *hrs*, if rounded to one significant figure. The most important source of the variation of coupling rates a and b is the diurnal variation **cite**. The peak times during which there is an appreciable exchange of matter between the free atmosphere and the ABL happens in the morning and evening hours, with significantly less exchange during the night, and even during the day. Therefore, the dominant period on which variation occurs is of the order of ~ 12 *hrs*, or in our units this is *τ* ≈ 4, and the the amplitude would be somewhere in between 0.5 and 1. The middle panel of Fig. 36 shows a peak in the speed of the front in the GL at *τ* ~ 10, or about 1 day, but the amplitude is likely larger than 0.5. Thus, the dominant period of the variation lies below the resonance peak. However, the numbers we provide here are a rough estimate. The most uncertain figure is the value of *δ*. It appears that the only accurate statement that we can make at this point is that the period diurnal variation is within an order of magnitude of the period at which a resonance peak occurs. This is a tantalizing observation.

Now, in the absence of modulation, the speed of the front, given *a = b* = 0.1 would be ≈ 0.945 in these units. In comparison to this, the amplitude of the speed variation in the presence of modulation is actually very modes. For example, we see from the middle panel of Fig. 36 that the amplitude of the speed variation is at most 0.15, which is about 16% of the unmodulated speed. We will now look at modulations of other parameters to explore if the effect will be larger.

### D.2 Varying the growth rate

The original model in physical variables is given by Eqs. (3)–(4). In those variables, the speed of the downwind front is given by

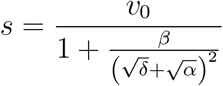

in the absence of diffusion. Here the advective speed is a cosntant, *v = v*_0_. The same model in dimensionless variables is given by Eqs. (5)–(6). In these variables, the dimensionless speed (i.e. speed in units of advective speed *v*_0_) is given by

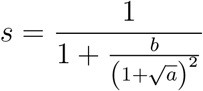

The growth rate *δ* becomes 1 in dimensionless variables. We chose to explore the role of modulating this growth rate, by replacing the growth term by 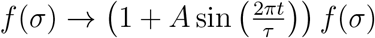. Then, in the adiabatic limit *τ* → 0, the front speed would approach

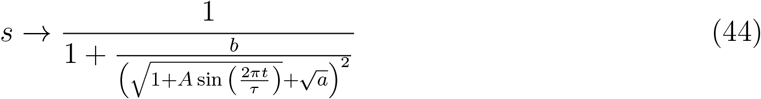

(consider the dimensionless model as a special case of the physical model with *v*_0_ = 1, *α* = *a, β* = *b*, and 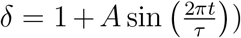. The corresponding min and max values would then be given by

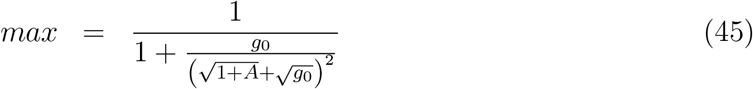

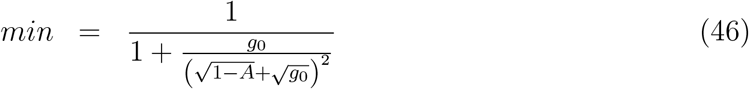

We summarize the resonance behavior with this kind of variation in the next graph. Here, the resonance peak is much higher. Evidently, the variation of the growth rate can achieve much more impressive variations of the front speed, in relation to the unmodulated value – even with A not approaching 1.

**Figure 37:**
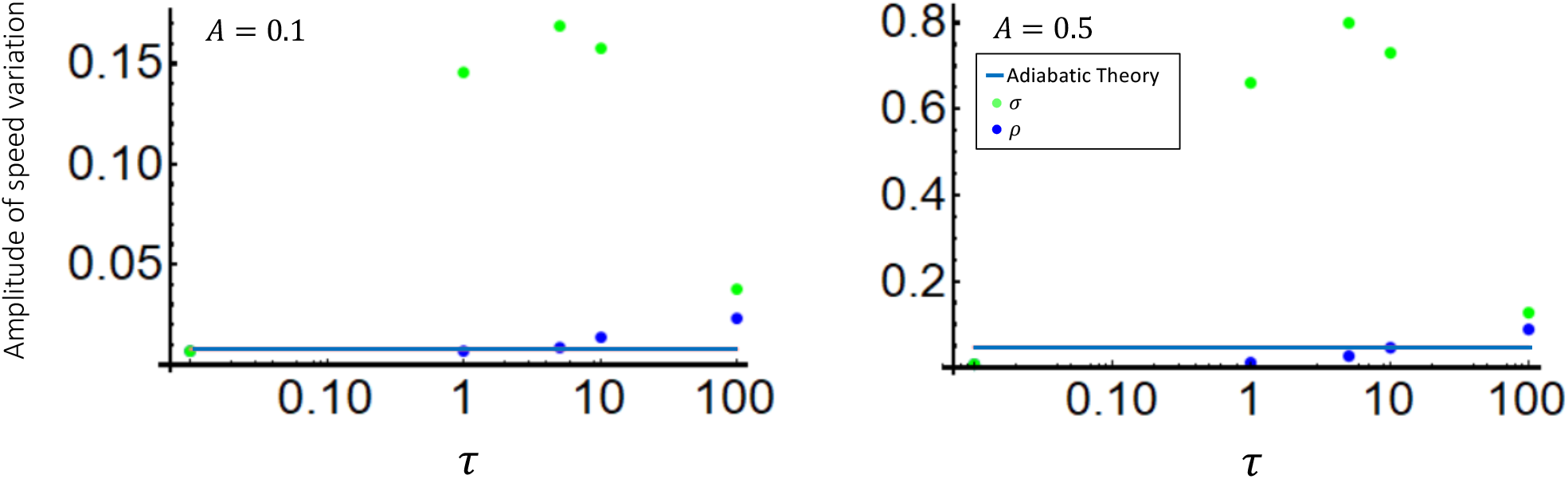
(Color online) Amplitude of speed variation vs. the period *τ* of the variation of the growth rate. Some green and blue dots coincide.

### D.3 Varying the wind speed

We have also varied the wind speed. The adiabatic predictions for this variation is

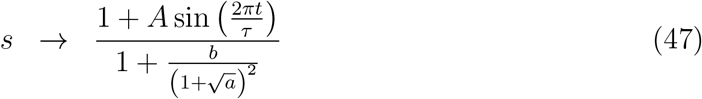

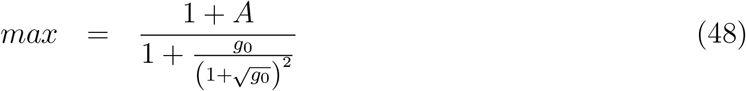

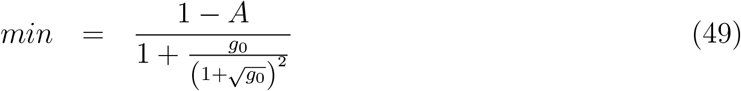

The following is a plot of the front speed variation vs. *τ* at two values of A.

**Figure 38:**
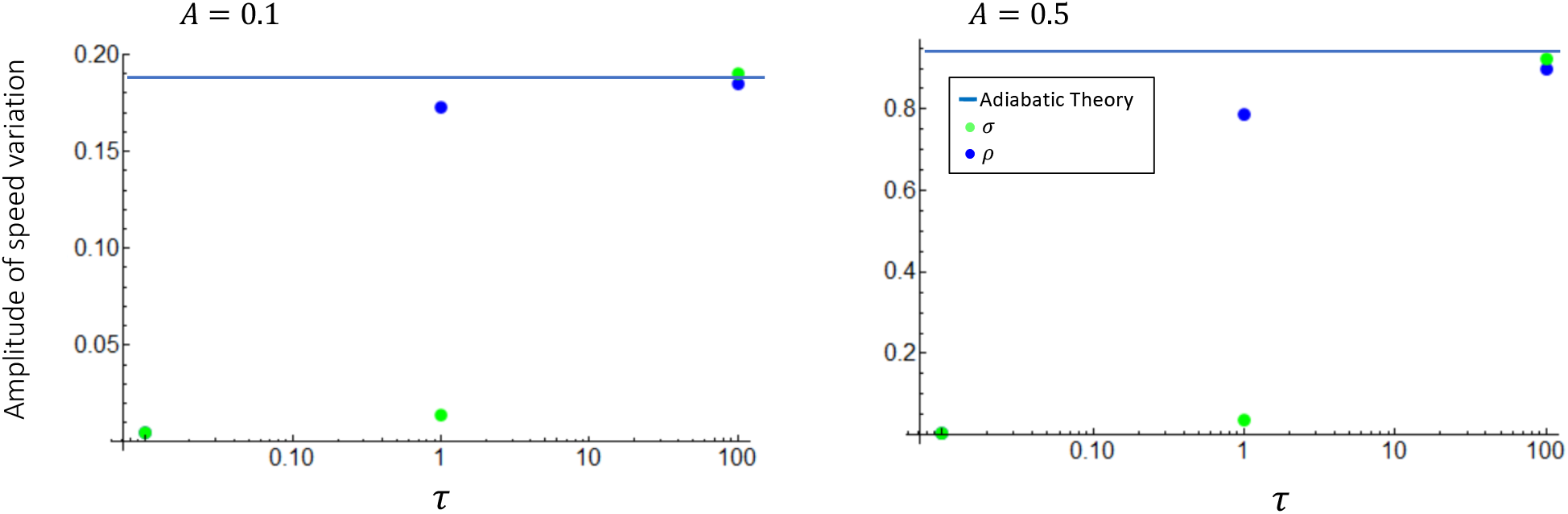
(Color online) Amplitude of speed variation vs. the period *τ* of the variation of the growth rate. Some green and blue dots coincide.

Here there is no resonance curve. Instead, for both *ρ* and *σ* there is a transition from a “low-τ” regime to an adiabatic “high-*τ*” regime. The transition takes place around different *τ* for ρ and *σ* – the transition for *ρ* occurs at lower *τ*.

## Footnotes

1 We used “FindRoot” in Mathematica. In doing this numerical root finding, we supplied the value 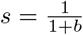 as an initial guess at any 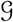. Sometimes, it helped to also supply the other value, 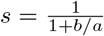 to create a smooth plot of s vs. 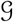. Using the “Solve” feature of Mathematica, we were also able to find all the roots, but only one of them satisfied both limits, and this is the root that was found with “FindRoot” with the aforementioned initial guesses.

